# Extensive opsin gene expansion and non-cerebral origin of the minimalist eye in a model tardigrade

**DOI:** 10.64898/2026.04.15.718617

**Authors:** Soumi Dutta, Vladimir Gross, Lars Hering, Mercedes Klein, Silja Flenner, Imke Greving, Elena Longo, Georg Mayer

## Abstract

Panarthropod vision exhibits extraordinary morphological and functional diversity, yet the sensory biology of tardigrades—microscopic extremophiles renowned for their resilience—remains poorly understood. In the model tardigrade *Hypsibius exemplaris*, we uncover an unprecedented expansion of opsin genes, with over 100 paralogs constituting the largest known opsin repertoire in any animal. Paradoxically, the visual system is structurally minimalist: a paired, inverse pigment-cup ocellus embedded within the brain lobes, forming a single-pixel, dual-receptor organ. Integrating genomic, phylogenetic, molecular expression, and ultrastructural analyses, we show that directional vision relies on a single rhabdomeric opsin (He-R-Opsin-V), localized to microvilli of the rhabdomeric cell. A ciliary photoreceptor with a lamellated cilium co-expresses two ciliary opsins (He-C-Opsin-1 and -2), suggesting non-visual light detection. These and other non-visual opsins are differentially expressed in the brain, gut, storage cells, and peripheral tissues, implicating them in circadian regulation, neuromodulation, ecdysis, digestion, and environmental sensing. Crucially, the eye is an internalized epidermal vesicle, not a cerebral derivative, challenging long-standing assumptions about its evolutionary origin. These findings reveal how extreme miniaturization drives sensory system simplification in visual organs while enabling parallel evolutionary innovation in non-visual photoreception. This study establishes a new paradigm for sensory evolution in microscale animals.

## Introduction

Photoreception is one of the most ancient and diverse sensory modalities in animals^1^. Within Panarthropoda—comprising Onychophora (velvet worms), Tardigrada (water bears), and Arthropoda—visual systems range from the simple ocelli of onychophorans^2,3^ to the complex compound eyes of insects^4–6^. Resolving the evolutionary origins and diversification of these systems requires understanding both the molecular basis of photoreception, particularly the opsin photopigments^7–11^, and the developmental and evolutionary origins of the photoreceptive organs themselves^3,9,12,13^.

Opsins are light-sensitive, G-protein-coupled receptors that mediate a wide array of light-driven processes, including image formation, circadian entrainment, and photoperiodic regulation^14^. Traditionally, photoreceptor cells are classified into two types based on their membrane origin: rhabdomeric photoreceptors (derived from microvilli and expressing r-opsins) and ciliary photoreceptors (derived from modified cilia and expressing c-opsins)^15^. With advances in genomic and transcriptomic data, opsins have been further classified into four major lineages: canonical visual opsins (including r-opsins, c-opsins, and bathyopsins), chaopsins, xenopsins, and tetraopsins (encompassing neuropsins, Go-opsins, and RGR/retinochromes)^16^. Despite extensive study in many animal lineages, opsin-based photoreception remains poorly understood in miniaturized, morphologically simplified organisms such as tardigrades.

Tardigrades occupy a pivotal phylogenetic position for reconstructing panarthropod evolution^17–21^. This question is complicated, however, by their body plan having been profoundly shaped by extreme miniaturization, involving cell number reduction, loss of organ systems, and the simplification or loss of major body regions^22–24^. How this constraint has influenced sensory evolution remains unclear. Current knowledge is marked by contradictions: while behavioral studies confirm light responsiveness^25–27^, estimates of opsin gene numbers in the model species^28,29^ *Hypsibius exemplaris* vary widely^10,11^, obscuring the true extent of its photoreceptive capabilities. Moreover, the evolutionary origin of the tardigrade eye remains controversial: some studies^30^ suggest an epidermal origin, consistent with other panarthropods, while others propose a unique intracerebral position and cerebral derivation^31,32^, distinct from their closest relatives.

Here, we resolve these long-standing ambiguities by integrating genomic and transcriptomic analyses with cDNA and gDNA cloning, comprehensive phylogenetic reconstruction, *in situ* hybridization, immunohistochemistry, and three-dimensional (3D) ultrastructural analysis of the visual system in *H. exemplaris*. We uncover a striking paradox: despite extreme morphological simplification of the eye, the genome harbors an unprecedented expansion of opsin genes—over 100 paralogs, the largest known repertoire in any animal. We demonstrate that the tardigrade eye functions as a single-pixel, directional sensor integrated into the brain. Crucially, we provide definitive cytological evidence that, despite its intracerebral location, the eye is a derived epidermal vesicle. These findings reveal that miniaturization in tardigrades has driven a functional decoupling of vision and non-visual photoreception, resulting in a highly reduced ocular structure alongside a complex, widespread system of extra-ocular and extra-cerebral light sensing.

## Results

### Massive expansion of the non-visual opsin repertoire

To resolve the complexity of the *H. exemplaris* opsin repertoire, we combined transcriptomic cDNA with genomic DNA cloning. This approach revealed a gene count substantially higher than previous estimates (Table 1). While confirming a single visual rhabdomeric opsin (*He*-*r-opsin-v*) and three ciliary opsins (*He-c-opsin-1* to *-3*; Supplementary Table 1), we uncovered a dramatic expansion of non-visual lineages. Of 56 analyzed *He-r-opsin-nv* transcripts, 48 are unique sequences, 17 (∼35L%) of which contain premature stop codons, indicating likely pseudogenes. The proportion of pseudogenes is even higher in neuropsins: among 119 cloned *He-neuropsin* homologs, 113 transcripts are unique, 59 (∼52L%) of which contain premature stops, primarily due to alternative splicing, especially exon 2 skipping. Nevertheless, 31 full-length functional *He-r-opsin-nv* and 54 *He-neuropsin* transcripts remain, representing one of the most extensive opsin expansions in any animal. Genomic cloning confirmed this diversity, revealing multiple distinct *He*-*neuropsin* loci and supporting their status as genuine paralogs rather than post-transcriptional isoforms.

**Table 1.**
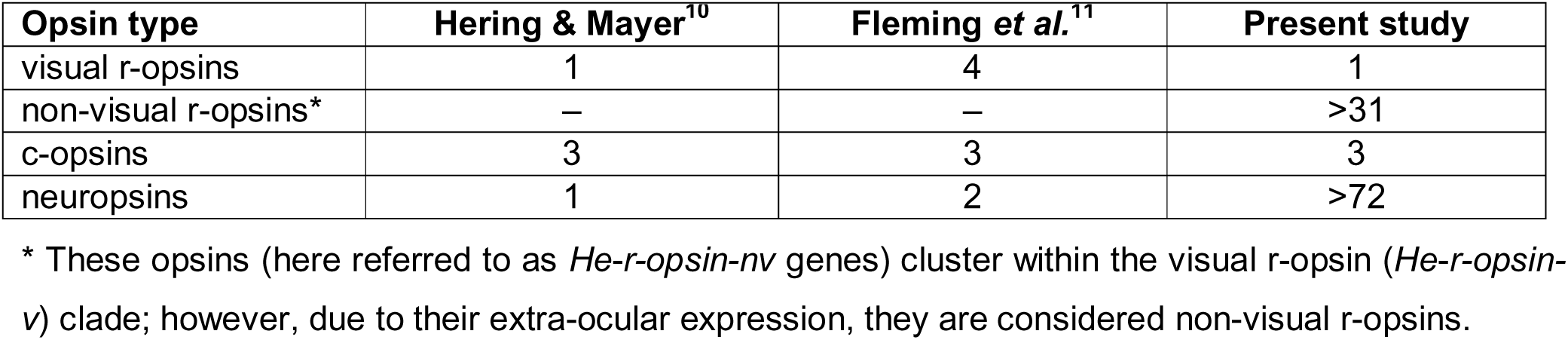
Previously reported and newly identified opsins in *H. exemplaris*. Note that the actual number of non-visual r-opsins and neuropsins is likely substantially higher than currently documented.

| Opsin type | Hering & Mayer<sup>10</sup> | Fleming *et al.*<sup>11</sup> | Present study |
| --- | --- | --- | --- |
| visual r-opsins | 1 | 4 | 1 |
| non-visual r-opsins* | – | – | >31 |
| c-opsins | 3 | 3 | 3 |
| neuropsins | 1 | 2 | >72 |

\* These opsins (here referred to as *He-r-opsin-nv* genes) cluster within the visual r-opsin (*He-r-opsin-v*) clade; however, due to their extra-ocular expression, they are considered non-visual r-opsins.

Phylogenetic analysis of seven-transmembrane domains places *He*-*r-opsin-v* within a clade sister to the visual r-opsin of *Ramazzottius varieornatus*, confirming its orthology as the primary visual photopigment (Fig. 1). In contrast, the expanded *He*-*r-opsin-nv* lineage—designated non-visual r-opsins—forms a sister group to the eutardigrade visual opsin clade and clusters with onychophoran visual opsins (*onychopsin* clade). *He*-*r-opsin-nv* sequences segregate into two well-supported subclades (*He-r-opsin-nv-a*, *-b*) (Fig. 1; Supplementary Fig. 1), indicating early lineage-specific diversification. Similarly, *neuropsin* paralogs form two distinct subclades (*He-neuropsin-a*, *-b*) (Fig. 2a) that cluster with broad protostome and deuterostome *neuropsin/opsin-5* sequences (Supplementary Fig. 1), suggesting deep evolutionary origins. The three c-opsins (*He-c-opsin-1* to *-3*) form a tardigrade-specific group within a clade of panarthropod canonical c-opsins (Fig. 1; Supplementary Fig. 1). Notably, no additional *He-c-opsin* paralogs were detected in the genome or via cloning, indicating that the massive gene expansion is specific to *He-r-opsin-nv* and *He-neuropsin* lineages.

**Fig. 1.**
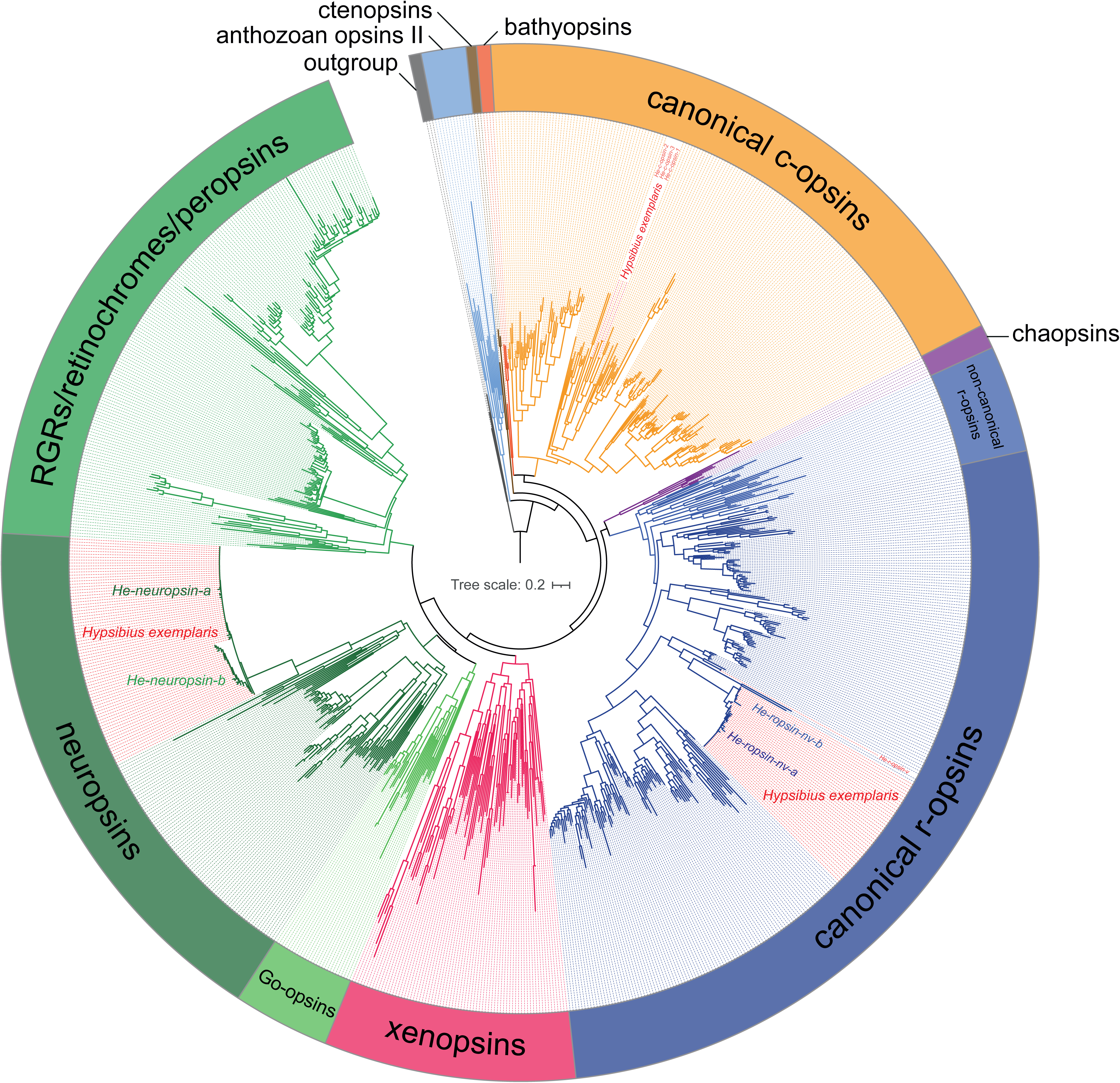
Phylogeny and orthology of metazoan opsins reveal extensive diversification in *Hypsibius exemplaris*. Maximum-likelihood tree of 906 metazoan opsins inferred under the GTR+G model, with nomenclature following Ramirez *et al.*^16^. *H. exemplaris* opsin genes are highlighted in red. The tree resolves a single visual r-opsin (*He*-*r-opsin-v*), a lineage of non-visual r-opsin paralogs (He-*r-opsin-nv-a/b*), three ciliary opsins (*He*-*c-opsin-1* to *-3*), and a large expansion of *He-neuropsin-a/b* homologs. Scale bar indicates substitutions per site. The full tree with bootstrap support values and accession numbers is provided in Supplementary Fig. 1.

**Fig. 2.**
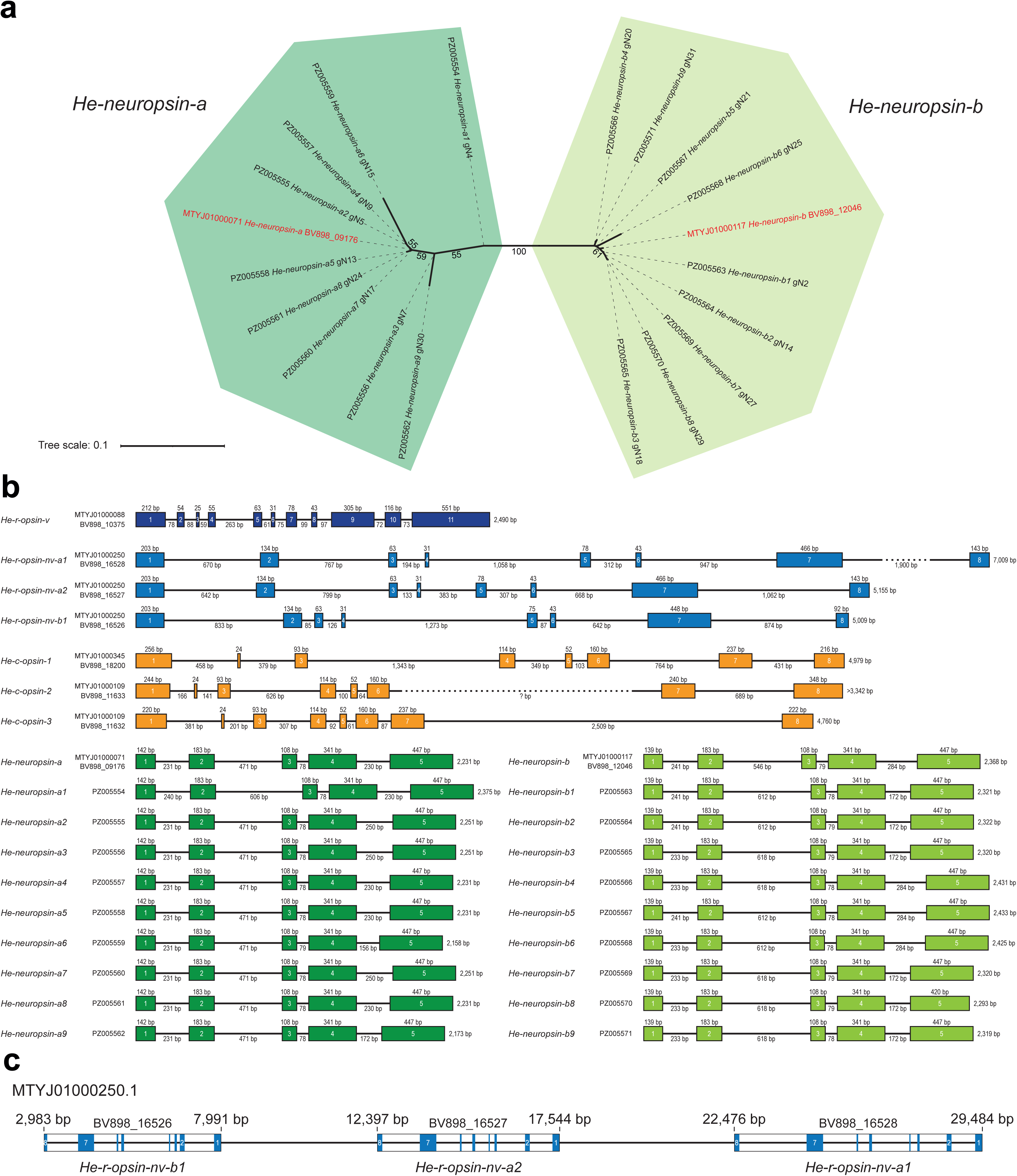
*He-neuropsin* phylogeny and genomic structure of opsin genes in *Hypsibius exemplaris*. (**a**) Unrooted phylogeny of full-length *He-neuropsin* sequences (including introns) cloned from genomic DNA, together with two sequences retrieved from the published genome of *H. exemplaris* (highlighted in red). Note the clear separation into two well-supported clades (*He-neuropsin-a* and *- b*). Scale bar indicates substitutions per site. (**b**) Diagrams illustrating the structure and length of opsin gene loci in *H. exemplaris*. Exons are depicted as colored rectangles and introns as solid black lines, with their respective lengths indicated. Alphanumeric labels correspond to GenBank accession numbers and locus tag IDs from the published genome. (**c**) Diagram of genomic scaffold250 (GenBank accession MTYJ01000250.1) illustrating the tandem arrangement of *He-r-opsin-nv* genes. Numbers indicate the positions of the genes on the scaffold. Exons are numbered and depicted as blue rectangles. Note the reverse orientation of all three genes, reflecting their localization on the complementary strand.

Genomic structure analysis revealed 5–11 exons across opsin genes (Fig. 2b). *He-r-opsin-v* has 11 short exons; *He-r-opsin-nv* and *He-c-opsin* genes have eight; and all *He-neuropsin* genes have five. Despite similar exon numbers and lengths, intron lengths vary dramatically in the expanded *He-r-opsin-nv* and *He-neuropsin* groups (Fig. 2b). All genomic clones differ in sequence (Fig. 2a,b), suggesting at least 20 distinct genomic origins. Notably, three *He-r-opsin-nv* loci (BV898_16526, BV898_16527, BV898_16528) are tandemly arranged on scaffold250 (Fig. 2c), indicating gene duplication-driven expansion (Fig. 2c).

Functional conservation is preserved across all opsins. All *H. exemplaris* opsins retain the retinal-binding lysine (K296) and the **NPxxY** motif (absent in neuropsins) in transmembrane helix 7 (Supplementary Fig. 2). He-R-Opsin-V retains the canonical **HPK** motif in the fourth cytoplasmic loop—critical for G_q_-coupled phototransduction^33,34^, while He-R-Opsin-NV paralogs exhibit a derived **HQR** motif. The three c-opsins retain the canonical **NxQ** motif for G_t_ activation^35^. Notably, He-Neuropsin proteins contain threonine (**T90**) at the position corresponding to **G90** of the bovine rhodopsin, and tyrosine (**Y91**) instead of lysine (**K91**) in human OPN5—both key determinants of UV sensitivity^10,36^ (Supplementary Fig. 3), suggesting potential roles in UV detection.

### Organization of a single-pixel, inverse pigment-cup eye

The visual organs of *H. exemplaris* are paired, inverse pigment-cup ocelli (volume ∼90 µm^3^), embedded within the outer brain lobes (Fig. 3a, b; Supplementary Fig. 4). 3D ultrastructural reconstruction reveals a minimalist architecture of exactly four cells per eye: a pigment cell, a rhabdomeric (retinula) cell, a ciliary cell, and a support cell (Fig. 3c–g; Supplementary Figs 5–8). None of these cells project axons; instead, five axon-bearing neurons lie medially adjacent, serving as the primary efferent pathway (Supplementary Fig. 9).

**Fig. 3.**
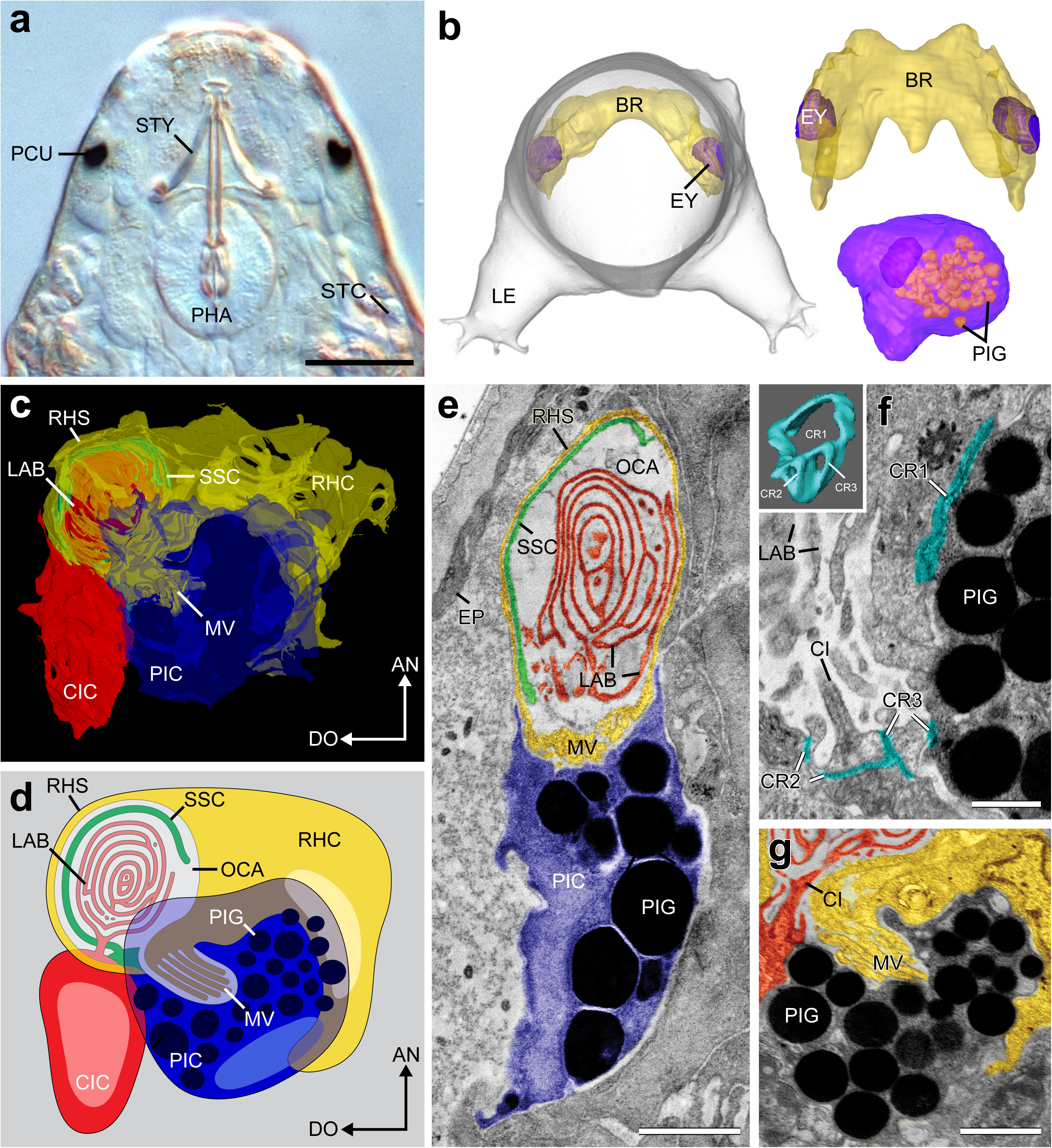
Ultrastructure and 3D organization of the *H. exemplaris* eye. (**a**) Differential interference contrast micrograph of the head in dorsal view. (**b**) 3D reconstruction from X-ray nanoCT showing the anterior body (frontal view), brain (dorsal view), and eye shape. Eyes are large relative to pigment spots in (a). (**c**) 3D model based on serial STEM-in-SEM ultrathin sections (anterior: up, dorsal: left). All four ocular cells are present; axons are absent. (**d**) Simplified schematic of eye organization; same depiction as in (c) (**e–g**) Pseudocolored STEM-in-SEM micrographs: pigment cell (blue), rhabdomeric cell (yellow), ciliary cell (red), support cell (green), Cytotardin junctions (turquoise). (**e**) Horizontal section showing the optic cavity, lamellar labyrinth, and support sheath (anterior: up, lateral: left. (**f**) Sagittal view of the cilium and lamellar labyrinth; inset: 3D reconstruction of Cytotardin rings (CR1–CR3) connecting all four eye cells. (**g**) Sagittal view illustrating rhabdomeric microvilli extending into the pigment cup (anterior: up, dorsal: left). Abbreviations: AN, anterior; BR, brain; CI, cilium; CIC, ciliary cell; DO, dorsal; EP, epidermis; EY, eye; LAB, lamellar labyrinth; LE, leg; MV, microvilli; OCA, optic cavity; PCU, pigment cup; PHA, pharynx; PIC, pigment cell; PIG, pigment granules; RHC, rhabdomeric cell; RHS, rhabdomeric sheath; SSC, sheath of support cell; STC, storage cell; STY, stylets. Scale bars: 20Lµm (**a**), 1Lµm (**e, g**), 500Lnm (**f**).

The pigment cell exhibits strong horseradish peroxidase (HRP) immunoreactivity (Fig. 4a–g) and contains ∼50 electron-dense granules, forming a cup that shields photoreceptive structures (Fig. 3a–g). The cup opens antero-dorso-laterally, accommodating the microvilli of the rhabdomeric cell (Fig. 3c–g; Supplementary Fig. 8). Crucially, apical junctions between ocular cells contain Cytotardin (Figs 3f, 4a–c; Supplementary Video 1)—a tardigrade-specific cytoplasmic intermediate filament protein that stabilizes epidermal and foregut tissues^37^. Three belt-like Cytotardin rings (CR1–CR3) provide definitive molecular evidence for an epidermal rather than neural origin of the ocular epithelium.

**Fig. 4.**
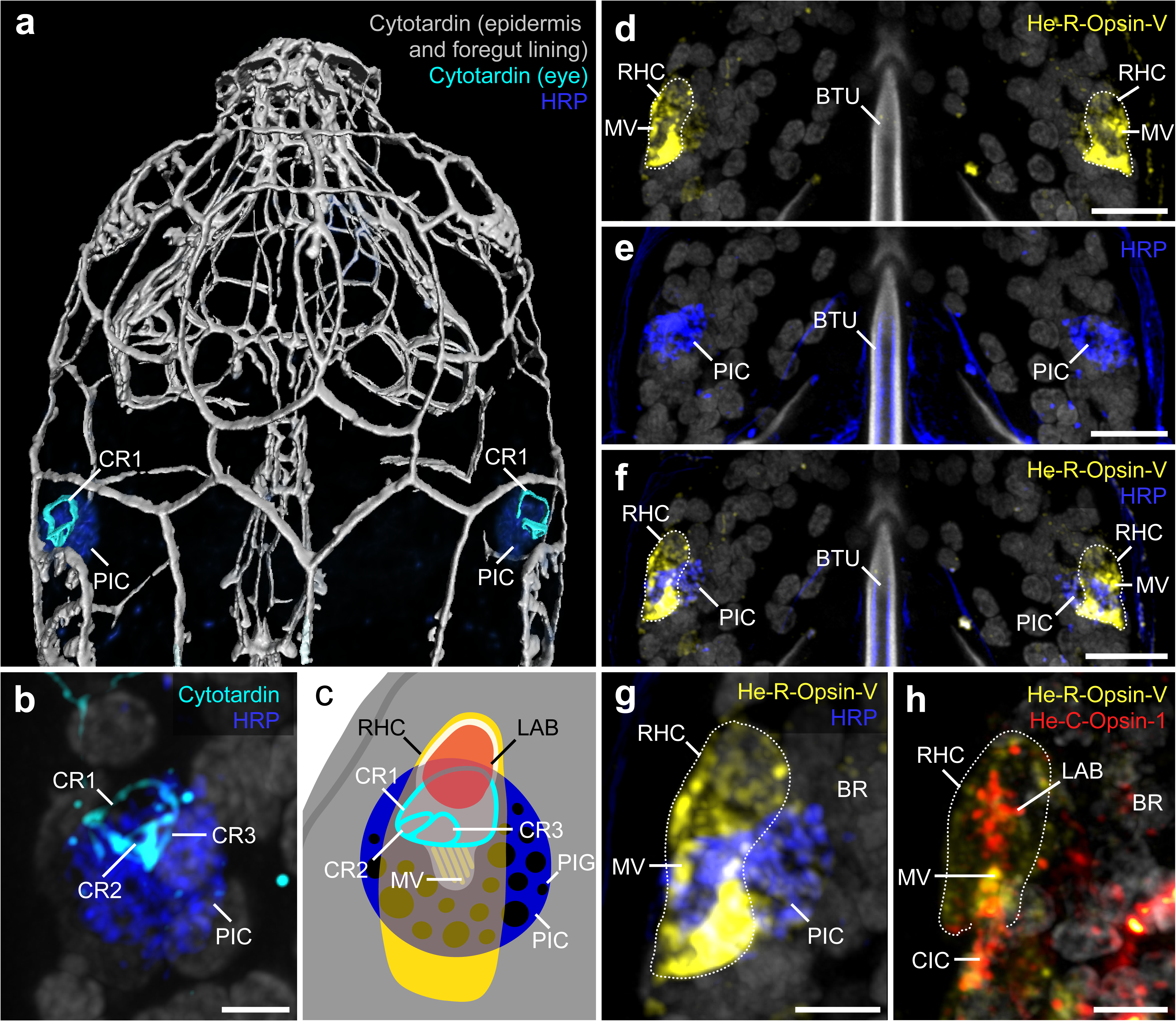
Subcellular localization of opsins and structural proteins in the *H. exemplaris* eye. Immunolabeling of Cytotardin (turquoise), HRP (blue), He-R-Opsin- V (yellow), and He-C-Opsin-1 (red); DNA in gray. Anterior: up. (**a**) Dorsal view showing Cytotardin rings (turquoise) at the pigment cell interface; Cytotardin also present in epidermal and foregut lining cells (volume-rendered gray meshwork). (**b**) High-magnification view of one large and two smaller Cytotardin rings. (**c**) Schematic of structures in (a, b, d–h). (**d–f**) He-R-Opsin-V (yellow) localized to rhabdomeric microvilli adjacent to the pigment cell (blue). (**g**) Close-up of left eye from (f). (**h**) Double labeling showing He-R-Opsin-V in rhabdomeric microvilli (yellow) and He-C-Opsin-1 in the lamellar labyrinth (red). Abbreviations: BR, brain; BTU, buccal tube; CIC, ciliary cell; CR1–CR3, Cytotardin rings; LAB, lamellar labyrinth; MV, microvilli; PIC, pigment cell; RHC, rhabdomeric cell. Scale bars: 2Lµm (**b, g, h**), 5Lµm (**d–f**).

The eye exhibits a dual-receptor organization. The rhabdomeric cell, with a flat nucleus, projects microvilli into the pigment cup groove (Fig. 3c–g; Supplementary Fig. 8). Simultaneously, the ciliary cell extends a specialized cilium into the optic cavity, anterior to the pigment shield. This cilium expands into a lamellar labyrinth—structurally analogous to the outer segment of vertebrate photoreceptors—laterally enclosed by a “support sheath” derived from the support cell (Fig. 3c–g; Supplementary Figs 5–8).

### Spatial distribution of visual and non-visual opsins

To map functional specialization, we combined Hybridization Chain Reaction–Fluorescence *In Situ* Hybridization (HCR-FISH) with immunohistochemistry using custom-made antibodies against He-R-Opsin-V, He-C-Opsin-1, and He-Neuropsin. As expected for a visual photopigment, He-R-Opsin-V is strictly confined to the rhabdomeric photoreceptor cell of the eye (Fig. 4c–h; Supplementary Fig. 10; Supplementary Video 2). However, the eye is not merely a rhabdomeric sensor: both *c-opsin-1* and *c-opsin-2* are co-expressed in the adjacent ciliary photoreceptor cell (Fig. 5a,b; Supplementary Figs 11, 12; Supplementary Videos 3–5). Notably, He-C-Opsin-1 protein localizes specifically to the distal lamellar labyrinth within the optic cavity (Fig. 4c,h), suggesting a role in light capture or signal amplification.

**Fig. 5.**
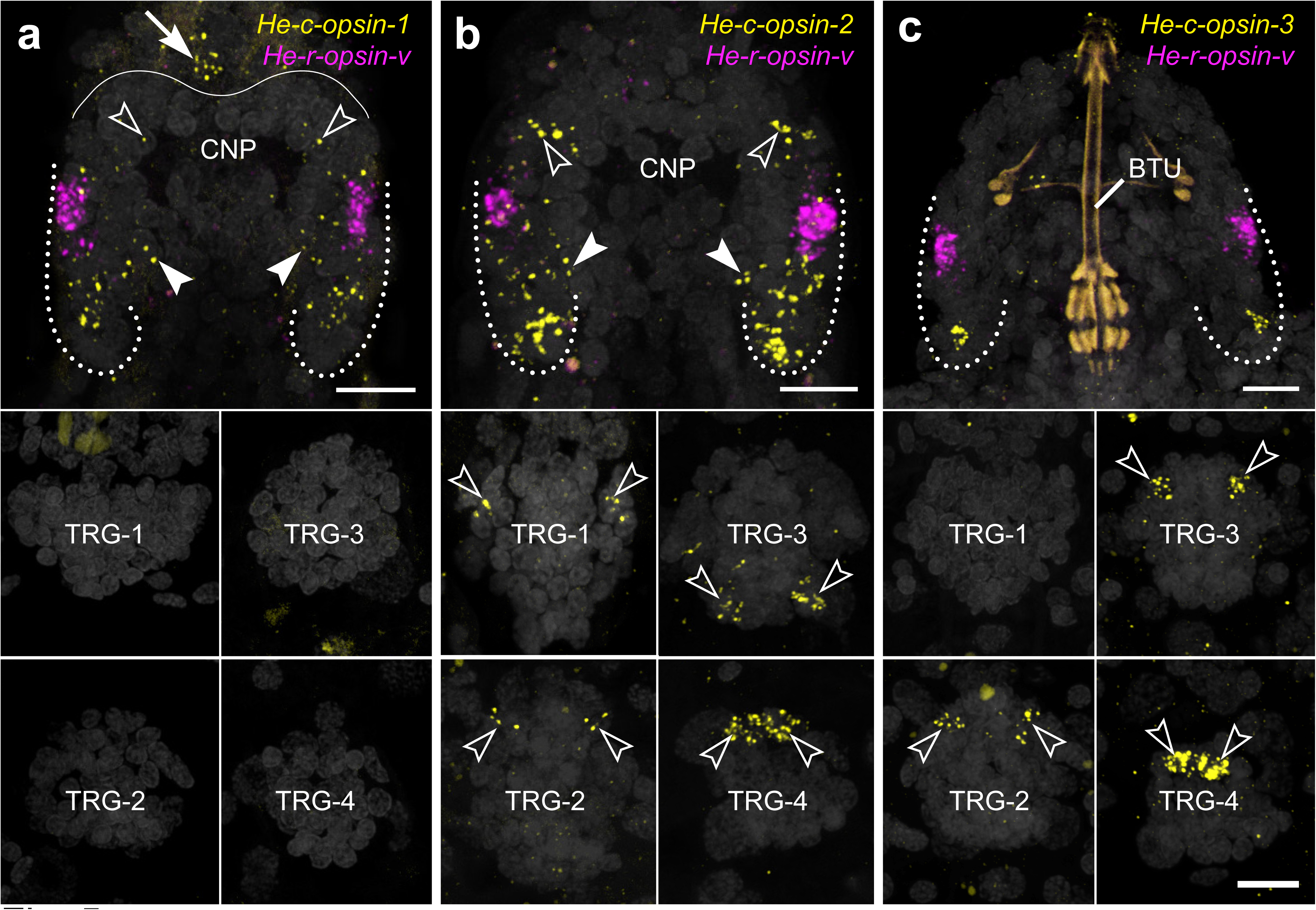
Expression of ciliary opsins in the central nervous system of *H. exemplaris*. HCR-FISH of *He*-*c-opsin-1*, *He*-*c-opsin-2*, *He*-*c-opsin-3* (yellow), and *He*-*r-opsin-v* (magenta); DNA in gray. Anterior: up. Brain: dorsal view; trunk ganglia: ventral view. (**a**) *He*-*c-opsin-1* is expressed in the posterior outer brain lobe (dotted line), inner brain lobe (filled arrowheads), anterior brain (open arrowheads), and dorsomedial extracerebral tissue (arrow). No expression in trunk ganglia. A weak, with *r-opsin-v* non-overlapping signal in the eye is observed. (**b**) *He*-*c-opsin-2* expression recapitulates *He*-*c-opsin-1* in the eye and brain with higher intensity. Notably absent in dorsomedial extracerebral tissue but shows additional expression in anterior neurons of ganglia 1, 2, and 4, and posterior neurons of ganglion 3 (open arrowheads). (**c**) *He*-*c-opsin-3* is restricted to single neurons in the posterior outer brain lobe and anterior neurons of ganglia 2–4. No expression detected in ganglion 1. Abbreviations: BTU, buccal tube; CNP, central brain neuropil; TRG-1–4, first to fourth ventral trunk ganglia. Scale bars: 5Lµm.

Beyond the eye, c-opsins exhibit broad functional pleiotropy. In addition to their co-expression in the ciliary cell, *He*-*c-opsin-1* and *-2* are co-localized in anterior and posterior brain clusters, with *He*-*c-opsin-2* showing stronger signal (Fig. 5a,b; Supplementary Figs 10–12). *He*-*c-opsin-2* is further expressed in all four trunk ganglia (Fig. 5b), with additional labeling in 1–2 cells per leg—likely peripheral leg ganglia^38^—within the first and third trunk segments, but not in the second or fourth (Supplementary Fig. 13). Notably, *He*-*c-opsin-1* and *He c-opsin-2* are also detected outside the central nervous system (CNS), with *He-c-opsin-1* expressed in a dorsomedian cluster anterior to the brain (Fig. 5a; Supplementary Figs 10, 12; Supplementary Video 3), and both opsins co-expressed in two pairs of ventrolateral clusters on each side of the head (Supplementary Videos 4, 5). These extracerebral clusters may represent gland-like perioral structures previously documented in other tardigrade species^39,40^. *He*-*c-opsin-3* is restricted to a posterior neuron in each outer brain lobe, an anteromedian cluster (in two specimens) located anterior to the central brain neuropil, and an anterior pair of neurons in the second to fourth trunk ganglia (Fig. 5c; Supplementary Fig. 12; Supplementary Video 6). Co-expression with *He*-*c-opsin-2* is observed in the second and fourth ganglia, suggesting functional synergy (Fig. 5b,c).

The expanded *He*-*r-opsin-nv* and *He*-*neuropsin* lineages—detected via pooled HCR-FISH probes (Supplementary Fig. 14)—show no expression in the eye but are localized to the CNS (Fig. 6a–e; Supplementary Figs 10, 14–16). Both are expressed (but not co-expressed) in a medial brain region adjacent to the eye and in posterior outer brain lobes (Supplementary Videos 7, 8), indicating non-visual roles in central sensory integration. *He*-*r-opsin-nv* is further expressed in one or two cells in the inner lobe (Supplementary Video 7), but not in trunk ganglia (Supplementary Fig. 15). *He*-*neuropsin* additionally marks a specific pair of posterior neurons in the first trunk ganglion, but is absent from the second to fourth (Fig. 6e; Supplementary Fig. 16).

**Fig. 6.**
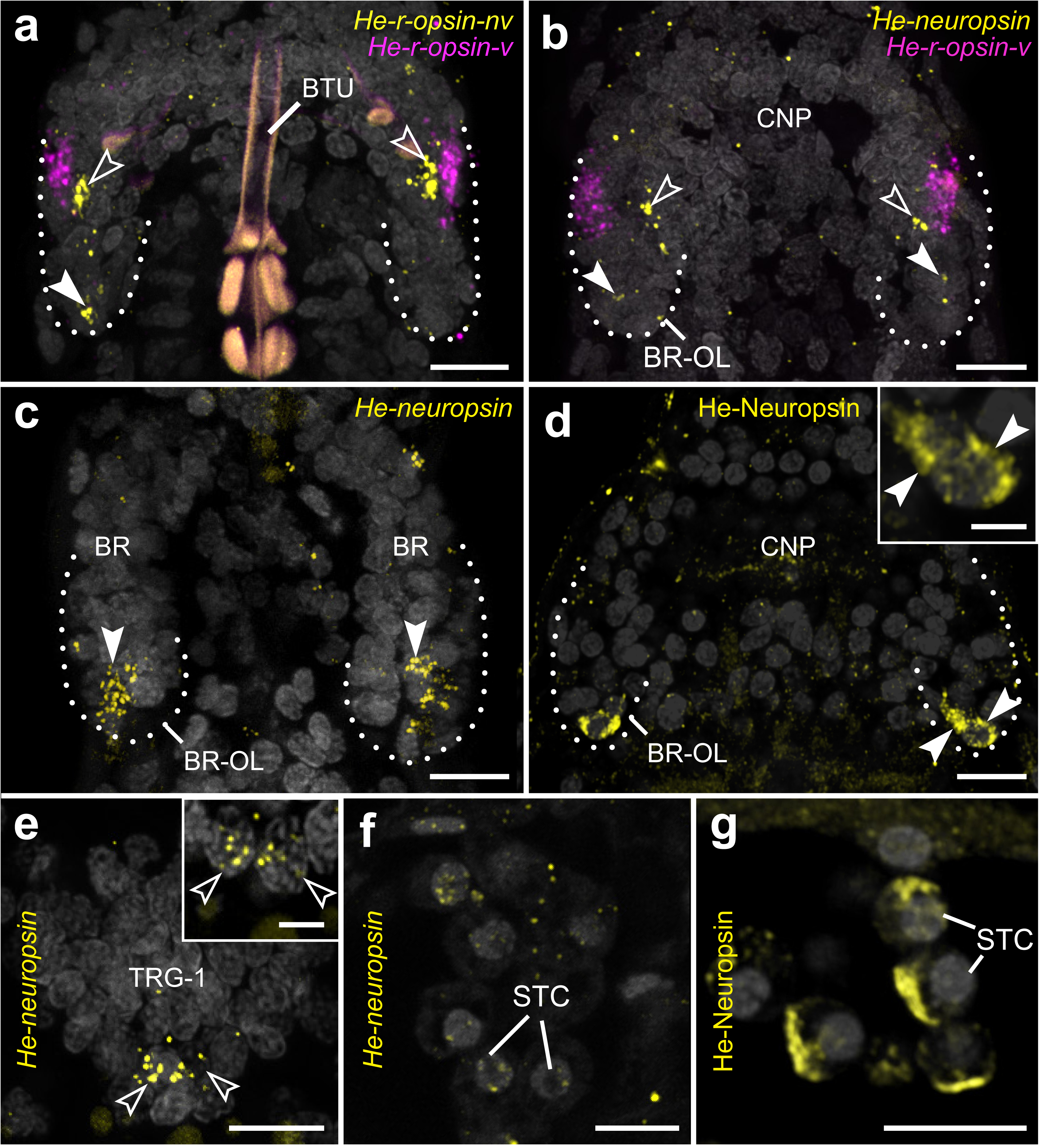
Expression of non-visual r-opsins and neuropsins in *H. exemplaris*. HCR-FISH (mRNA) of *He-r-opsin-nv* and *He-neuropsin* (both yellow) and *He-r-opsin-v* (magenta); immunolabeling (protein) of He-Neuropsin (yellow in **d**, **g**), DNA in gray. Anterior: up. Brain: dorsal view; trunk ganglion: ventral view. (**a**) *He*-*r-opsin-nv* expressed adjacent to *He*-*r-opsin-v* in the eye (open arrowheads) and in posterior outer brain lobe (filled arrowhead). (**b**) *He*-*neuropsin* mRNA in anterior (open arrowheads) and posterior (filled arrowheads) outer brain lobe regions. (**c, d**) *He*-*neuropsin* mRNA and protein, respectively, in posterior outer brain lobe (filled arrowheads); inset: magnified posterior region. (**e**) *He*-*neuropsin* mRNA in posterior region of first trunk ganglion (open arrowheads); inset: magnified posterior region showing one pair of neuropsin immunoreactive cells. (**f, g**) *He*-*neuropsin* mRNA and protein localized to storage cells, with potential protein enrichment in a polarized membrane stack (possibly Golgi). Abbreviations: BR, brain; BR-IL, inner brain lobe; BR-OL, outer brain lobe; BTU, buccal tube; CNP, central brain neuropil; STC, storage cell; TRG-1, first trunk ganglion. Scale bars: 5Lµm (**a–g**), 2Lµm (inset in **d, e**).

Most strikingly, *He*-*neuropsin* is expressed in a subset of midgut and storage cells (Fig. 6f,g; Supplementary Fig. 17). Immunolabeling reveals the He-Neuropsin protein localizes to a polarized membrane stack—possibly the Golgi apparatus—in storage cells (Fig. 6g). This subcellular localization, combined with the proliferative activity of midgut and storage cells^41,42^, points to a novel non-visual function of opsins in light-dependent regulation of metabolism, energy storage, or cell cycle dynamics.

To validate specificity, HCR-FISH negative controls (no DNA probes, hairpin amplifiers only) showed no signal in brain, midgut, or storage cells (Supplementary Fig. 17). Blocking peptide assays abolished immunolabeling in tissues previously positive (Supplementary Fig. 18). Autofluorescence in calcified and sclerotized foregut structures (cuticle, stylets, pharyngeal placoids^43^) and external cuticle (including claws) fluctuates across the molting cycle (Figs 4, 5; Supplementary Figs 10,14, 18)—intense before molting, absent after sheding^44^—confirming it is intrinsic autofluorescence, not specific labeling.

## Discussion

### A molecular toolkit for non-visual photoreception in a miniature animal

The opsin repertoire of *Hypsibius exemplaris* reveals an unprecedented degree of gene family expansion and functional diversification, fundamentally challenging prior estimates of only five to nine opsin genes^10,11^. By combining genomic and transcriptomic cloning with phylogenetic reconstruction, we estimate a minimum of 107 opsin genes, with the potential for further expansion as additional clones are sequenced from the respective genomic loci. This represents the largest known opsin repertoire in any animal, surpassing even the crustacean *Daphnia pulex*, which encodes 48 opsins^45^.

This expansion is driven by one canonical visual r-opsin (*He-r-opsin-v*), three ciliary opsins (*He-c-opsin-1* to *-3*), and at least 31 non-visual r-opsins (*He-r-opsin-nv*) and 72 *He-neuropsin* homologs. The *He-r-opsin-nv* lineage clusters within the visual r-opsin clade but exhibits hallmarks of tandem duplication—occurring adjacent on scaffold250—and accelerated sequence divergence, including a high proportion of pseudogenes. These features suggest neofunctionalization and a transition from visual to non-visual roles. Similarly, both *He-r-opsin-nv* and *He-neuropsin* paralogs form two distinct subclades (*a/b*), indicating an ancient duplication followed by lineage-specific diversification—possibly driven by adaptive pressures in a microscale, environmentally variable niche.

Functional implications are evident in spatial expression patterns. While He-R-Opsin-V is strictly confined to the rhabdomeric photoreceptor—confirming its role as the primary visual photopigment^10^—the *He*-*r-opsin-nv* and *He*-*neuropsin* paralogs are expressed in non-ocular brain regions (Fig. 7a), including a medial cluster adjacent to each eye and a posterior brain cluster per hemisphere. These regions are anatomically proximate to neurons expressing Pigment-Dispersing Factor (PDF) neuropeptides^46^, suggesting a role in non-visual light input to the circadian system—akin to Rhodopsin 7 in *Drosophila*^47^ or extra-retinal opsins in vertrebrates^14^. Notably, both gene lineages are enriched in eggs compared to adults^11^, hinting at roles in embryogenesis or photomorphogenesis, but this warrants further study.

**Fig. 7.**
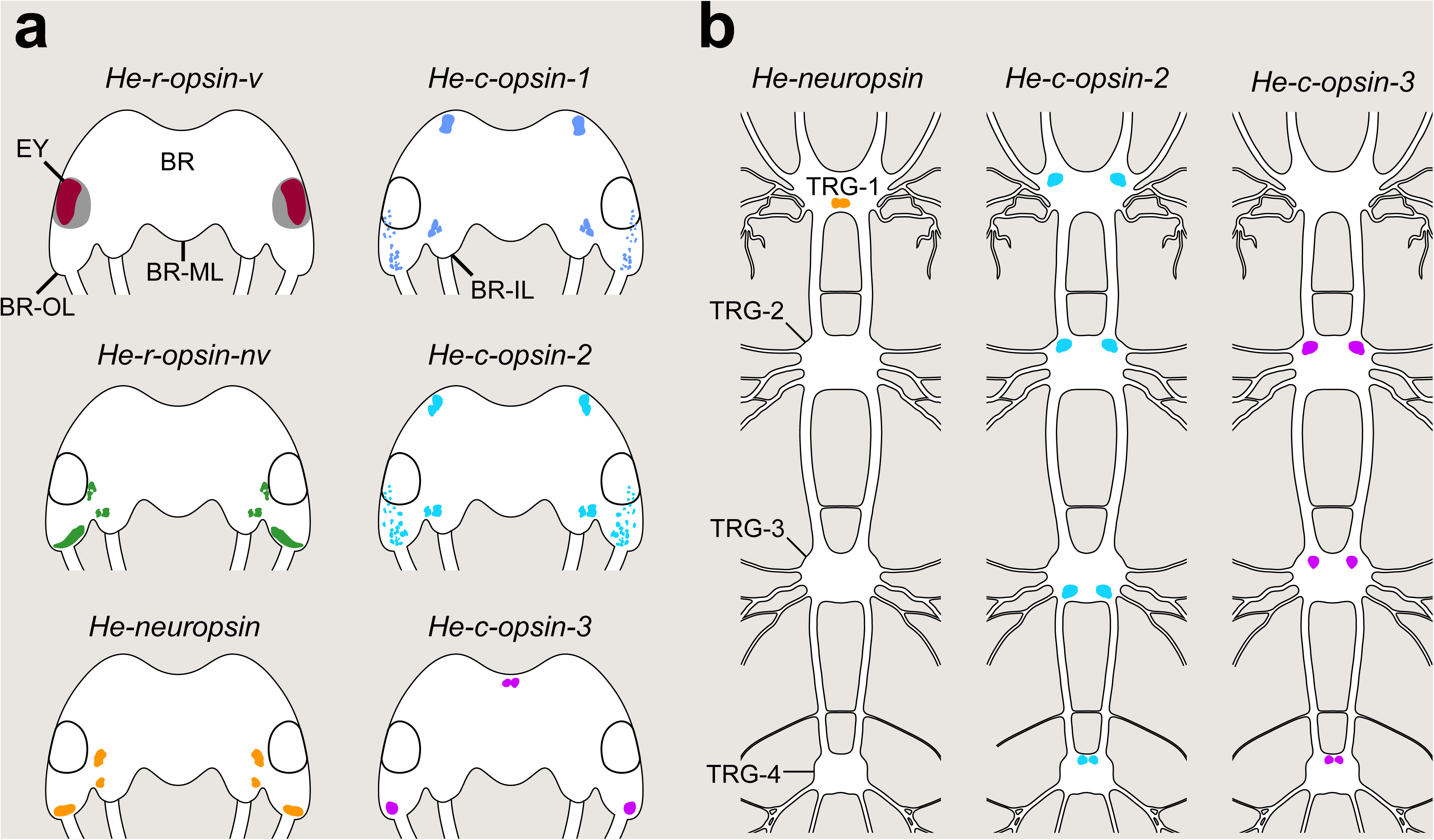
Integrated map of opsin expression across the *H. exemplaris* nervous system. (**a, b**) Dorsal (a) and ventral (b) views of the central nervous system. Anterior: up. All opsin classes are expressed in the brain, but *He*-*r-opsin-v*, *He*-*r- opsin-nv*, and *He*-*c-opsin-1* are absent from trunk ganglia. *He*-*c-opsin-3* is not expressed in TRG-1, and *He*-*neuropsin* is absent from TRG-2–4. The spatial segregation of opsins suggests a functional specialization. Abbreviations: BR, brain; BR-IL, inner brain lobe; BR-ML, median brain lobe; BR-OL, outer brain lobe; EY, eye; TRG-1–4, first to fourth ventral trunk ganglia.

Ciliary opsins exhibit even greater functional complexity. *He*-*c-opsin-1* and *-2* co-localize in the ciliary photoreceptor and multiple brain clusters, while *He*-*c-opsin-3* is restricted to posterolateral brain regions (Fig. 7a). *He*-*c-opsin-2* is expressed in all four trunk ganglia and *He-c-opsin-3* in trunk segments 2–4 (Fig. 7b), suggesting roles in peripheral sensory integration and segmental coordination. Intriguingly, *He*-*c-opsin-1* and *-2* are detected in extracerebral tissues: an unpaired dorsomedian cluster and two paired ventrolateral structures, morphologically consistent with perioral glands previously described in other tardigrade species^39,40^. This spatial association raises the possibility of a role in molting, including (stylet) cuticle shedding and remodeling—processes regulated by light and circadian cues in other panarthropods^48–50^.

Most strikingly, *He*-*neuropsin* is expressed in midgut and storage cells—tissues not previously linked to opsin function. We speculate that in storage cells, immunolabeling may localize the protein to polarized membrane stacks—possibly the Golgi apparatus—suggesting a role in light-dependent regulation of metabolism, energy storage, or cell cycle dynamics. This aligns with the widespread expression of the PDF receptor in tardigrade tissues^46^. In *Drosophila*, PDF receptor activation modulates circadian rhythms via cAMP signaling^51^. Given that midgut and storage cells of *H. exemplaris* undergo active proliferation^41,42^ and may co-express *He*-*neuropsin* and PDF receptors^46^, we propose a novel system for light-dependent control of systemic physiology. Motif analysis reveals that all *He-neuropsin* genes encode threonine at position 90 (**T90**) and tyrosine at position 91 (**Y91**)—a well-established site for UV sensitivity^10,36^. This represents a radical extension of opsin function beyond traditional neural and sensory roles^52–56^.

Together, these findings reveal a sophisticated, multi-layered photoreceptive system in *H. exemplaris*. Visual and non-visual opsins (r-, c-, and neuropsins) are spread across neural and peripheral tissues to mediate vision, circadian regulation, ecdysis, digestion, and metabolism, demonstrating how massive gene family expansion enables complex sensory integration even in a miniature body plan.

### Single-pixel, stereoscopic “vision”: a minimalist solution to directional light detection

The eyes of *H. exemplaris* exemplify extreme miniaturization and functional optimization. Ultrastructural and 3D reconstructions confirm paired, inverse pigment-cup ocelli embedded in the outer brain lobes—structurally equivalent to those of other tardigrades^30–32^. Despite their simplicity, they exhibit a tetracellular organization: a pigment cell, a rhabdomeric photoreceptor, a ciliary photoreceptor, and a support cell. None project axons, indicating secondary sensory function.

The pigment cell contains electron-dense granules, whose biochemical composition remains unknown^31^, and exhibits strong horseradish peroxidase (HRP) immunoreactivity. The consistent antero-dorso-lateral orientation of each pigment cup suggests a structural adaptation for stereoscopic or directional light detection—a hallmark of directional photoreceptors^57^. This is supported by documented light-oriented behaviors in tardigrades^25–27^, which likely rely on interocular signal differences.

The rhabdomeric cell expresses a single opsin, He-R-Opsin-V, localized mainly to microvilli—indicating monochromatic vision. With only one photoreceptor and photopigment, the eye functions as a photon counter, not an image-forming organ. Spatial resolution is fundamentally limited^57^, and the system likely operates under high photon shot noise, especially in dim light^58^. Yet, when combined with stereoscopic input from two eyes, it can support behaviorally relevant phototaxis, potentially tuned to blue-green wavelengths, aligning with peak sensitivity in onychophorans^59^.

The ciliary photoreceptor bears a distal lamellar labyrinth—structurally analogous to vertebrate outer segments^60^—suggesting convergent evolution for maximizing photopigment density. Co-expression of He-C-Opsin-1 and -2 raises the possibility of spectral broadening^61^ or enhanced sensitivity^62^, though their spectral properties remain unknown. Crucially, the labyrinth lies outside the pigment cup, arguing against a role in directional vision. Instead, it likely functions as a non-visual photodetector—perhaps for UV avoidance or environmental monitoring—mirroring extra-retinal photoreception in other animals^63,64^.

Thus, the *H. exemplaris* eye is not an image-forming organ but a highly specialized, single-pixel, stereoscopic photoreceptor system optimized for directional light detection. This minimalist architecture reflects the constraints of miniaturization, which has profoundly shaped all organ systems in tardigrades^24,65^.

### Epidermal origin of tardigrade eyes: a re-evaluation of evolutionary homology

The origin of tardigrade eyes has long been debated. Early interpretations proposed an epidermal origin based on integumental association^30^, while later studies on *Milnesium tardigradum* argued for a cerebral origin due to an intracerebral location^31,32^. Our ultrastructural data resolve this controversy: in *H. exemplaris*, the eyes are fully integrated into the brain but lack morphological continuity with the epidermis.

Key features distinguish ocular cells from neurons: absence of axons, epithelial organization, and expression of Cytotardin—a tardigrade-specific cytoplasmic intermediate filament protein that stabilizes epidermal and foregut tissues^37^. The presence of three belt-like Cytotardin rings at the apical junctions of the four ocular cells further reinforces their epithelial identity. These structures, previously misclassified as desmosomes or zonulae adhaerentes^66–68^, are functionally analogous to adherens junctions in other bilaterians but are molecularly distinct, highlighting the evolutionary innovation of the tardigrade skin.

This combination of features—epithelial organization, absence of axons, and expression of epidermis-specific structural proteins—strongly supports an epidermal origin of the tardigrade eye, despite its intracerebral integration. This aligns with developmental patterns in other panarthropods: in onychophorans, eye *anlagen* invaginate directly from the embryonic epidermis^3,9^; in arthropods, both median ocelli and compound eyes derive from non-neurogenic ectoderm^12,13^. These similarities suggest the ancestral panarthropod eye was an epithelial vesicle.

The inverse cup morphology of tardigrade eyes—where the rhabdomeric microvilli face away from incoming light—appears derived, likely resulting from extreme miniaturization and reduction of a multicellular pigment layer to a single cell^3,65,69^. This contrasts with the ancestral everse configuration in onychophorans and arthropods, which retain the epidermal origin but differ in morphological elaboration^3,5^. The pit-like invagination in arthropods may represent a heterochronic arrest at a transient developmental stage—recapitulated in onychophoran development^9,70^—rather than a fundamental divergence.

In sum, our findings support an epidermal origin for the tardigrade eye, reconciling morphological and molecular data. This re-interpretation underscores the importance of molecular markers—such as Cytotardin—in resolving deep evolutionary homologies.

## Conclusion

This study reveals that *Hypsibius exemplaris* possesses a remarkably diversified opsin toolkit, with paralogs spread out across neural and peripheral tissues to mediate visual and non-visual light sensing, potentially including circadian and physiological roles. The eye, though structurally minimalist, functions as a stereoscopic, monochromatic photoreceptor optimized for directional light detection. Despite its intracerebral position, its molecular and cellular architecture—epithelial organization, absence of axons, and expression of epidermis-specific proteins—strongly supports an epidermal origin, in line with tardigrades’ closest relatives in the panarthropod lineage. Together, these findings reshape our understanding of sensory evolution in microscopic animals and highlight the power of integrating ultrastructure, genomics, and molecular expression analysis to uncover hidden complexity in the smallest of animals.

## Methods

### Rearing of animals

Specimens of the tardigrade *Hypsibius exemplaris* Galsiorek, Stec, Morek & Michalczyk, 2018^71^ (Eutardigrada, Parachela, Hypsibiidae; Sciento, Whitefield, Manchester, UK; strain Sciento Z151^72^) were maintained in plastic Petri dishes and fed with unicellular algae *Chlorococcum* sp. under a 12:12 h light–dark cycle at 21 °C as described elsewhere^23,46^.

### Amplification and cloning of opsins from cDNA and gDNA

Total RNA was extracted from several hundred *H. exemplaris* specimens using TRIzol reagent (Invitrogen, Carlsbad, CA, USA) and purified with the RNeasy MinElute Cleanup Kit (Qiagen, Hilden, Germany). The species is parthenogenetic, resulting in extremely low levels of genetic heterozygosity among individuals, as offspring are produced clonally from unfertilized eggs^28,73^. First-strand cDNA was synthesized with random hexamer primers and SuperScript IV reverse transcriptase (Invitrogen), following the manufacturer’s protocol. Genomic DNA (gDNA) was isolated from several hundred *H. exemplaris* specimens using the DNeasy Blood & Tissue Kit (Qiagen). Complete coding sequences of the nine reported *H. exemplaris* opsins^11^ were amplified from first-strand cDNA as well as from genomic DNA (*He-neuropsin-a/b*) using gene-specific primers (Supplementary Table 2) and cloned into the pJET1.2/blunt vector with the CloneJET PCR Cloning Kit (Thermo Fisher Scientific, Waltham, MA, USA). Colony PCR products were verified by Sanger sequencing and sequences were deposited in GenBank under accession numbers PX981454–PX981555 (cDNA), PX981588–PX981664 (pseudogenes), and PZ005554–PZ005571 (gDNA).

### Phylogenetic analyses

Previously identified opsin sequences from tardigrades (*H. exemplaris* and *Ramazzottius varieornatus*)^10,11^, together with all known opsin sequences from onychophorans^8,59,74^, were aligned with the metazoan opsin dataset of Ramirez *et al.*^16^ using MAFFT v7.505^75^ with the G-INS-i algorithm. The alignment was masked with Noisy v1.5.12^76^ to remove homoplastic and randomly aligned positions (parameters: -cutoff=0.80 -distance=HAMMING -missing=N -nogap=0 -noconstant=0 -ordering=nnet -shuffles=20000 -smooth=1 -seqtype=P). A maximum-likelihood tree was inferred using the Pthreads version of RAxML v8.2.12^77^, conducting ten independent runs to identify the best-scoring topology under the empirical LG+G4+F substitution model (-f a option) obtained using ModelTest-NG^78^ implemented in raxmlGUI v2.0^79^. Node support was estimated from one hundred rapid bootstrap pseudoreplicates. Full *He-neuropsin* sequences (including introns) cloned from genomic DNA as well as two sequences obtained from the publicly available genome of *H. exemplaris* (GenBank accession: MTYJ00000000.1) were aligned likewise without subsequent masking. The best maximum-likelihood tree was inferred from 100 independent runs (GTR+G) and 1,000 thorough bootstrap pseudoreplicates. The resulting trees were visualized with iTOL v6.6^80^ and edited in Adobe Illustrator CS5.1 (Adobe, San Jose, CA, USA).

### Scanning electron microscopy

Specimens of *H. exemplaris* were prepared and examined as described previously^23^, with the following modification: imaging was performed using a Hitachi S-4000 field-emission scanning electron microscope (Hitachi High-Technologies Europe GmbH, Krefeld, Germany) operated at 7 kV instead of 5 kV.

### Ultrathin sectioning and scanning transmission electron microscopy in a scanning electron microscope (STEM-in-SEM)

Specimens were asphyxiated and fixed in 4 % paraformaldehyde and 1 % glutaraldehyde in 0.1 mol L^−1^ Sørensen sodium phosphate buffer (27.6 g L^−1^ NaH_2_PO_4_⋅H_2_O and 28.4 g L^−1^ Na_2_HPO_4_, mixed at a ratio of 23:77 and diluted 1:1 with double-distilled water; pH 7.4). After 1 h of fixation, samples were washed twice for 15 min in Sørensen buffer and postfixed in 1 % aqueous OsO_4_ for 1 h at room temperature. Specimens were rinsed overnight in distilled water and dehydrated through an ascending ethanol series (30 %, 50 %, 70 %, 90 %, 95 %; 10 min each) at room temperature, followed by two 30 min incubations in 100 % ethanol and two 30 min incubations in acetone.

Samples were infiltrated with Spurr’s resin (Spurr Low-Viscosity Embedding Kit, “standard” formulation; Sigma-Aldrich) by incubation in a 1:1 mixture of acetone and resin for 30 min, then twice in 100 % resin for 60 min each and overnight on a rotator at room temperature. After acetone evaporation, samples were incubated in fresh resin for 4–6 h before transfer into block-shaped silicone molds containing fresh resin. Five to six animals were manually aligned within each mold using fine needles to facilitate subsequent sectioning. Polymerization was carried out at 70 °C for 3 days.

Ultrathin sections (60–65 nm thick, based on silver–gold interference color) were cut with a diamond knife (Diatome AG, Biel, Switzerland) on a Reichert-Jung Ultracut E ultramicrotome (C. Reichert AG, Vienna, Austria). Ribbons of approximately ten sections were collected on Formvar^®^-coated, single-slot copper grids (Plano GmbH, Wetzlar, Germany). Sections were stained manually with uranyl acetate replacement stain (UAR-EMS; diluted 1:4 in 50 % methanol; Electron Microscopy Sciences, Hatfield, PA, USA) for 5 min, followed by lead citrate^81^ for 8 min at room temperature. Grids were rinsed three times in distilled water and rapidly dried by capillary contact with filter paper.

Samples were imaged using the STEM-in-SEM technique as described previously^23^, employing a custom-built STEM converter for ultrathin section imaging in a field-emission scanning electron microscope. Imaging was conducted using a Hitachi S-4000 field-emission SEM at 23–25 kV. Image series were aligned using a combination of semi-automated and manual landmark-based registration in the TrakEM2 plugin^82^ for the Fiji distribution of ImageJ2^83^.

### Nano-computed tomography (nanoCT)

Specimens were fixed in 4 % formaldehyde for 1 h at room temperature, rinsed five times in distilled water (10 min each), and dehydrated through an ascending ethanol series (30 %, 50 %, 70 %, 90 %, and 2 × 100 %) for 15–30 min each. To prevent shrinkage artifacts, samples were processed by critical-point drying using a Bal-Tec 030 Critical Point Dryer. Animals were attached by their posterior body region to a specimen holder (either a sewing needle or custom-made stub), with the head oriented upright to fit within the scan window.

X-ray nanoCT imaging was performed at beamline P05^84,85^ (DESY, Hamburg, Germany), which is operated by Helmholtz-Zentrum Hereon at 11 keV using a transmission X-ray microscope with Zernike phase contrast. The optical system included a beam shaper with 100 µm subfields and a Fresnel zone plate (250 µm diameter; outer zone width 50 nm). Phase rings were employed to generate Zernike contrast. Fly scans were conducted over 180° sample rotation at 0.223 seconds per rotation, yielding 1,589 projections with 0.5 seconds exposure each. Radiographs were acquired at 2048 × 2048 pixels, corresponding to an effective pixel size of 35.52 nm.

During reconstruction, the data were binned by factor 2, yielding a final voxel size of 71.04 nm, allowing analysis of the anterior body region from multiple orientations and planes. Due to the low contrast between soft tissues and sclerotized structures, phase-contrast effects were used to enhance visibility^86^. Variations in gray values enabled tissue differentiation, and labeled structures were segmented for 3D surface rendering and volumetric measurements.

### Custom antibody generation and immunohistochemistry

Custom polyclonal antibodies were generated against He-R-Opsin-V (ID No. 78-01-15; 0.8 mg mL^−1^), He-C-Opsin-1 (ID No. 2067-06-16; 0.75 mg mL^−1^), and He-Neuropsin (ID Nos. 30-10-17 and 30-11-17; 1.0 mg mL^−1^). Antibodies were purified by high-performance liquid chromatography (HPLC; Peptide Specialty Laboratories GmbH, Heidelberg, Germany). Specificities were validated by peptide competition assays following immunohistochemistry.

To obtain fully extended specimens, animals were asphyxiated in tap water at 60 °C for 30 min and immediately fixed in either 4 % formalin in 0.1 mol L^−1^ PBS (pH 7.4) or Zamboni fixative^87^ (4 % formalin and 7.5 % picric acid in 0.02 mol L^−1^ PBS) for 1–2 h at room temperature. During fixation, specimens were punctured with an electrolytically sharpened tungsten needle to enhance reagent and antibody penetration. After fixation, samples were washed repeatedly, including an overnight rinse. Next day, samples were treated with Collagenase/Dispase^®^ (Roche Diagnostics GmbH, Mannheim, Germany) and hyaluronidase (Sigma-Aldrich, St. Louis, MO, USA), each at 1 mg mL^−1^, for 10 min at 37 °C, followed by postfixation (15 min). For Zamboni-fixed specimens, additional washes were performed in NH_4_Cl (50 mmol L^−1^; 2 × 15 min) and glycine (0.1 mol L^−1^ in PBS; 30 min).

Blocking was performed in 10 % normal goat serum (NGS) in PBS-Tx for 1 h. Samples were then incubated with primary anti-opsin antibodies (Supplementary Table 3) in PBS containing 1 % NGS and 0.1 % NaN_3_ at 4 °C for 2–5 days. After washing in PBS-Tx, samples were incubated with secondary antibodies (Supplementary Table 4) under the same conditions for 2–4 days at 4 °C. Specimens were counterstained with DAPI (4’,6-diamidino-2-phenylindole; Carl Roth GmbH, Karlsruhe, Germany; 1–2 µg mL^−1^), mounted in ProLong™l Gold Antifade Reagent (Thermo Fisher Scientific), and sealed with nail polish. For double-labeling experiments, additional primary antibodies were applied, including rabbit anti-HRP (12 µg mL^−1^; Jackson ImmunoResearch) and custom-made guinea pig anti-Cytotardin^37^ (21 µg mL^−1^).

### Blocking peptide assay for antibody specificity tests

After optimizing antibody concentrations for immunolabeling (against He-R-Opsin-V, He-C-Opsin-1, and He-Neuropsin), specificity was validated using blocking peptide assays. Samples were fixed and processed as for the standard immunolabeling protocol. Synthetic peptides used for immunization were mixed with their corresponding antibodies at a 1:62.5 molar ratio (8 µg mL^−1^ antibody + 500 µg peptide in PBS containing 1 % NGS and 0.1 % NaN_3_) and incubated overnight at 4 °C with gentle rotation. The antibody–peptide mixtures were then applied to samples and incubated for 5 days at 4 °C. Following multiple washes in PBS-Tx, samples were incubated with secondary antibodies for 2–4 days at 4 °C, then counterstained and mounted as described in the immunohistochemistry section.

### Hybridization Chain Reaction – Fluorescence *In Situ* Hybridization (HCR-FISH)

Sets of 8–18 DNA split-initiator probes (each 52 nucleotides) specific to different *H. exemplaris* opsin mRNAs were designed and synthesized by Molecular Instruments (Los Angeles, CA, USA; Supplementary Table 5). Animal preparation and HCR-FISH procedures followed established protocols^46^. After HCR-FISH, specimens were counterstained with DAPI for 1 h and mounted in ProLong™l Gold Antifade Reagent (Thermo Fisher Scientific).

### Image acquisition and processing

Light micrographs were captured using a Zeiss Axio Imager M2 microscope (Carl Zeiss, Jena, Germany). Fluorescently labeled specimens were imaged with a Zeiss LSM 880 confocal laser-scanning microscope equipped with an Airyscan module. Raw Airyscan Z-stacks were processed with ZEN 2 (black/blue edition; Carl Zeiss Microscopy GmbH) using automatic filter strength. Confocal substacks, projections, and image adjustments were conducted in Fiji v.1.52^83,88^, and the LUT tool was used to apply false colors. 3D reconstructions were generated from nanoCT, STEM-in-SEM, and Airyscan datasets using segmentation and volume rendering in Amira 6.0.1 (Thermo Fisher Scientific). Final figures were composed in Adobe Photoshop 24.5.0 and Adobe Illustrator 27.6.1.

## Supporting information

Dutta_et_al_Supplementary_Materials

## Acknowledgements

We thank Sonja Fuhrmann and Sonja Kasten for assistance with immunohistochemistry and western blotting, Christine Martin and Niklas Metzendorf for support with confocal microscopy, and Sandra Treffkorn for providing the light micrograph of *H. exemplaris*. We are grateful to Milena Marie Grollmann for HRP immunolabeling, Lisa Epple for He-R-Opsin-Visual and HRP double labeling, Cathleen Sorgalla for He-R-Opsin-Visual and He-C-Opsin-1 double labeling, and Michael Klinger and Henry Jahn for help with 3D reconstructions. We thank Frank Smith and Mandy Game for sharing unpublished opsin expression data from juveniles. Stefan Fischer is acknowledged for assistance with preliminary electron microscopy, and Rick Hochberg for donating diamond knives. We acknowledge Helmholtz-Zentrum Hereon and DESY, members of the Helmholtz Association HGF, for the provision of the experimental infrastructure and beamtime at P05 at PETRA III for the proposals I-20170052 and I-20191380.

## Language editing support

The authors used ChatGPT (OpenAI) and DeepL (DeepL SE) to assist with language editing, grammar refinement, and stylistic improvements. All scientific content, data analysis, and interpretations were independently developed, reviewed, and verified by the authors. The use of these tools did not replace human judgment or scientific responsibility.

## Author contributions

G.M., L.H., V.G., and S.D. conceived and designed the study. S.D. performed cloning, HCR-FISH, immunolabeling, peptide competition assay, CLSM, data analysis, 3D reconstruction and designed the panels. V.G. performed CLSM, ultrathin sectioning, electron microscopy and analyzed nanoCT tomography and electron microscopy data. L.H. conducted transcriptomic, genomic, and phylogenetic analyses. M.K. prepared tardigrade samples for X-ray nanoCT tomography, performed ultrathin sectioning and carried out 3D reconstructions. I.G., S.F. and E.L. assisted with experiments at Beamline P05 and data reconstruction. V.G., S.D., and G.M. wrote the initial draft of the manuscript. All authors reviewed and approved the final version.

## Supplementary Figures

**Supplementary Fig. 1. Phylogenetic relationships and orthology of opsins in *Hypsibius exemplaris*.** Maximum-likelihood (ML) phylogenetic tree inferred under the LG+G4+F model of amino acid evolution, based on 906 opsin and opsin-related sequences. *H. exemplaris* opsin genes are highlighted in red. Bootstrap support values (>50 %) from 100 pseudoreplicates are shown at nodes. Scale bar indicates substitutions per site.

**Supplementary Fig. 2. Conserved motifs of visual, non-visual rhodopsin and ciliary opsin sequences in *H. exemplaris*.** Multiple sequence alignment of predicted key functional motifs (e.g., retinal-binding site **K296**, **NPxxY**, and tripeptide motif in the fourth cytoplasmic loop) across *H. exemplaris* opsin lineages (highlighted in light grey). Amino acid residue and motif comparison with bovine rhodopsin (GenBank accession: NM_001014890.2) revealed lysine at position **K296** and **NPxxY** in all *He-r-opsin-v*, *He-r-opsin-nv,* and *He-c-opsin* clones. Note the occurrence of canonical c-opsin fingerprint tripeptide **NxQ** in all *He-c-opsin*, the canonical r-opsin tripeptide **HPK** in *He-r-opsin-v* and a similar **HQR** motif in all *He-r-opsin-nv* sequences.

**Supplementary Fig. 3. Amino acid comparison in *H. exemplaris* neuropsin sequences.** Alignment of bovine rhodopsin (NM_001014890.2), human Opn5 (AY377391.1) and all *He-*neuropsin clones indicate the presence of threonine (**T**) at the position of 90 (**T90**) and tyrosine (**Y**) at the position of 91 instead of **K91** (highlighted in light grey) suggesting UV-sensitivity of the respective opsin. Amino acid numbering is based on the bovine rhodopsin.

**Supplementary Fig. 4. Virtual cross-sections from X-ray nanoCT of the *H. exemplaris* head.** Dorsal is up. (**a**) Anterior brain region showing the optic cavity at the junction of the pharynx and buccal tube. (**b**) Anterior boundary of the eye. Abbreviations: BR, brain; BR-IL, inner brain lobe; BR-OL, outer brain lobe; BTU, buccal tube; EY, eye; PHA, pharynx. Scale bars: 5 μm (**a, b**).

**Supplementary Fig. 5. Ultrastructure of the *H. exemplaris* eye.** STEM-in-SEM micrographs of ultrathin sections. (**a**) Horizontal section of the left eye (anterior: up, lateral: left). (**b**) Sagittal view of the ciliary cell’s lamellar labyrinth (inset: 3D reconstruction of Cytotardin rings CR1 to CR3). (**c**) Sagittal view showing spatial organization of ocular cells. Abbreviations: CI, cilium; CR1–CR3, Cytotardin rings; EP, epidermis; LAB, lamellar labyrinth; MV, microvilli; OCA, optic cavity; PIC, pigment cell; PIG, pigment granules; RHS, rhabdomeric sheath; SSC, sheath of support cell. Scale bars: 1 μm (**a, c**), 500 nm (**b**).

**Supplementary Fig. 6. Structural details of the lamellar labyrinth and rhabdomeric microvilli.** Series of horizontal sections (ventral to dorsal) through the optic cavity. Dorsal: top, lateral: left. (**a–j**) Note the irregular arrangement of lamellar membranes and the close apposition of rhabdomeric microvilli to the pigment cell. Abbreviations: LAB, lamellar labyrinth; MV, rhabdomeric microvilli; OCA, optic cavity; PIG, pigment granule; RHS, rhabdomeric sheath; SSC, support cell sheath. Scale bars: 500 nm (**a–j**).

**Supplementary Fig. 7. Deformation of the lamellar labyrinth in the *H. exemplaris* eye.** Serial horizontal sections showing displacement of the lamellar labyrinth toward the medial side of the optic cavity (**a**–**d**). Anterior: up, lateral: left. Abbreviations: LAB, lamellar labyrinth; MV, rhabdomeric microvilli; OCA, optic cavity; RHS, rhabdomeric sheath; SSC, support cell sheath. Scale bar: 500Lnm.

**Supplementary Fig. 8. Reconstruction of the *H. exemplaris* eye.** Sagittal (a), ventral (b), and dorsal (c) views of a 3D model based on serial ultrathin sections. (**a**) Spatial relationships among rhabdomeric, pigment, and ciliary cells (dorsal: left). (**b**) Ventral view of the lamellar labyrinth (blue) and support cell sheath (magenta), originating from a cilium dorsally (arrowhead) and extending laterally. (**c**) Tilted dorsal view confirming association of the lamellar labyrinth with the cilium (arrowhead). Abbreviations: CIC, ciliary cell; LAB, lamellar labyrinth; MV, rhabdomeric microvilli; NU, nuclei; PIC, pigment cell; RHC, rhabdomeric cell; SSC, support cell sheath.

**Supplementary Fig. 9. Axon-bearing neurons associated with the *H. exemplaris* eye.** Sagittal view. (**a**) Electron micrograph showing five axon-bearing neurons (pseudocolored) projecting anteriorly from the outer brain lobe. (**b**) 3D reconstruction of the same neurons and axons. Neurons are located medially to the eye. Abbreviations: AX-1–5, axon-bearing neurons 1 to 5; PIC, pigment cell; RHC, rhabdomeric cell. Scale bar: 2 µm (**a**).

**Supplementary Fig. 10. Spatial expression of opsin transcripts in the *H. exemplaris* head.** HCR-FISH confocal projections (anterior: up). Autofluorescence from the buccal apparatus is visible. *He*-*r-opsin-v* is restricted to the eyes (dotted outlines). Other opsins show distinct brain expression patterns, with partial co-expression of *He*-*c-opsin-1* and *He*-*c-opsin-2*. *He*-*r-opsin-nv* and *He*-*neuropsin* probes detect multiple paralogs. Arrow: extracerebral *He*-*c-opsin-1* in dorsomedian head region outside the brain. Abbreviations: BTU, buccal tube; CNP, central brain neuropil; RHC, rhabdomeric cell. Scale bars: 10 µm.

**Supplementary Fig. 11. Expression of ciliary opsins and visual r-opsin in the brain and eye.** Confocal substack projections in dorsal (a) and lateral (b, c) views. DNA counterstain: gray. (**a**) *He-c-opsin-1* in the eye and dorsolateral outer brain lobe. (**b**) *c-opsin-2* in the eye and outer brain lobe (arrowheads). (**c**) *He-c-opsin-3* in a posterior outer brain lobe cell (arrowhead). Abbreviation: BTU, buccal tube. Scale bars: 5 µm (**a–c**).

**Supplementary Fig. 12. Co-expression of *He-c-opsin-1* and *He-c-opsin-2*, in combination with or without *He-r-opsin-v*, and expression of *He-c-opsin-3* in the brain.** Dorsal view of a confocal substack. Anterior: up. DNA counterstain: gray. (**a**) Co-localization of *He-c-opsin-1* (yellow) and *c-opsin-2* (magenta) transcripts in the same brain clusters confirms shared expression domains. Open arrowhead indicates the adjacent ciliary photoreceptor cell, where both transcripts are co-expressed. (**b**) Co-expression of *He-c-opsin-1* (yellow) and *c-opsin-2* (magenta) in the adjacent ciliary photoreceptor cell of another specimen, using *He-r-opsin-v* (magenta) as a landmark. (**c**) Extracerebral expression of *He*-*c-opsin-1* in dorsomedian head region (arrow). (**d**) Expression of *He-c-opsin-3* in the anteromedian brain cluster (filled arrowhead). Abbreviations: BR-OL, outer brain lobe; BTU, buccal tube; CNP, central brain neuropil; RHC, rhabdomeric cell. Scale bars: 5 µm (**a–d**).

**Supplementary Fig. 13. *He-c-opsin-2* expression in the legs of *H. exemplaris*.** Lateral views of confocal substacks. Claws are autofluorescent. Anterior: left. DNA counterstain: gray. *He-c-opsin-2* is expressed in 1–2 cells (likely peripheral leg ganglia) in legs 1 and 3 (open arrowheads), but not in legs 2 and 4. Dotted lines: claw glands. Abbreviations: CLG-1–4, claw glands 1 to 4; CL, claw; LE-1–4, legs 1 to 4. Scale bars: 10 µm (**a–d**).

**Supplementary Fig. 14. Co-expression of non-visual r-opsin paralogs in the brain.** Double HCR-FISH of (a, b) *He-r-opsin-nv-a1* and *He-r-opsin-nv-a2*, and (c, d) *He-r-opsin-nv-b1* and *He-r-opsin-nv-a2*. Dorsal views, anterior: up. DNA counterstain: gray. (**a, c**) Co-localization in brain clusters; (**b, d**) close-ups of right hemisphere. Abbreviation: BTU, buccal tube. Scale bars: 5 µm (**a–d**).

**Supplementary Fig. 15. Absence of *He-r-opsin-nv* expression in ventral trunk ganglia.** Ventral views of confocal substacks. Anterior: up. DNA counterstain: gray. No signal detected in ventral trunk ganglia. Abbreviations: TRG-1–4, first to fourth trunk ganglia. Scale bars: 5 µm (**a–d**).

**Supplementary Fig. 16. *He*-*neuropsin* expression in the brain and absence in trunk ganglia.** Dorsal (a, b) and ventral (c–e) views. Anterior: up. DNA counterstain: gray. (**a, b**) *He*-*neuropsin* transcripts in medial brain regions near the eyes (open arrowheads) and in posterior outer/inner brain lobes (closed arrowheads). Inset (b): close-up of inner brain lobe. (**c–e**) No expression in second to fourth trunk ganglia. Abbreviations: BR, brain; BR-IL, inner brain lobe; BR-OL, outer brain lobe; CNP, central brain neuropil; TRG-2–4, second to fourth trunk ganglia. Scale bars: 5 µm (**a–e**), 2 µm (inset in **b**).

**Supplementary Fig. 17. *He-neuropsin* expression in the midgut of *H. exemplaris*.** Ventral (a, b) and lateral (c) views. Anterior: left. Dotted lines: esophagus and midgut outlines. (**a, b**) *He*-*neuropsin* transcripts in subsets of cells across anterior, middle, and posterior midgut. (**c**) Negative control (no probe) shows no signal. Abbreviations: ESO, esophagus; HE, head; HIG, hindgut; LE-1–4, legs 1 to 4; MIG, midgut; TRG-3, third trunk ganglion. Scale bars: 10 µm (**a–c**).

**Supplementary Fig. 18. Peptide competition assay confirming the specificity of immunolabeling for He-R-Opsin-V, He-C-Opsin-1, and He-Neuropsin in *H. exemplaris*.** Confocal substacks of the head in dorsal view. Anterior: up. DNA counterstain: gray. **(a–c)** Pre-incubation of each antibody with its corresponding peptide abolished signal in the eye and brain region. Abbreviations: BR, brain; BTU, buccal tube. Scale bars: 10 µm (**a–c**).

## Supplementary Tables

**Supplementary Table 1. Revised nomenclature for opsins in *H. exemplaris*.** Proposed standardized nomenclature for opsins identified in *H. exemplaris*, based on phylogenetic clustering, functional inference, and lineage-specific expansion. Includes canonical visual r-opsin (*He-r-opsin-v*), three ciliary opsins (*He-c-opsin-1* to *-3*), and expanded paralogous lineages: non-visual r-opsin (*He-r-opsin-nv*) and *He-neuropsin*. The naming reflects evolutionary relationships and functional divergence, facilitating cross-species comparisons.

**Supplementary Table 2. Gene-specific primers for amplification of *H. exemplaris* opsin transcripts.** List of forward and reverse primers used for RT-PCR amplification of opsin transcripts from first-strand cDNA. Primers were designed based on transcriptome sequences and validated for specificity. All amplicons were confirmed by Sanger sequencing and used for cloning, probe generation, and expression validation.

**Supplementary Table 3. Polyclonal antibodies against *H. exemplaris* opsins.** Polyclonal antibodies generated by Peptide Specialty Laboratories GmbH (Heidelberg, Germany) targeting unique peptide sequences of *H. exemplaris* opsins. Antibodies were validated by immunohistochemistry and peptide competition assays. Note: Due to high sequence similarity among *He-neuropsin* paralogs, cross-reactivity of the He-Neuropsin antibody with multiple proteins cannot be excluded.

**Supplementary Table 4. Secondary antibodies used in immunohistochemistry and peptide competition assays.** List of secondary antibodies used for signal amplification in immunohistochemistry and peptide competition assay analyses. Includes species-specific, fluorophore-conjugated antibodies (e.g., Alexa Fluor 488, 568). All were validated for minimal non-specific binding and used at optimized dilutions.

**Supplementary Table 5. HCR-FISH probe and amplifier details.** Comprehensive list of HCR-FISH probes and amplifier sequences used for *in situ* detection of opsin transcripts. Probes were designed to target unique regions of each opsin gene, with specificity confirmed by BLAST analysis. Amplifiers (B1–B4) were used to enhance signal sensitivity. All probes and amplifiers are compatible with multiplex detection and high-resolution confocal imaging.

## Supplementary Videos

**Supplementary Video 1. Cytotardin expression in the head of *H. exemplaris*.** High-resolution confocal stack reveals Cytotardin—a tardigrade-specific cytoplasmic lamin—in the epidermis, cephalic sensory fields, buccopharyngeal lining, and ocular epithelium. Anterior: up. Cytotardin forms apical belt-like junctions at the interface of the four photoreceptor cells in the eye, indicating an epithelial origin. The pigment cell is labeled with horseradish peroxidase antibody (HRP; blue). Cytotardin (turquoise) is absent from neural tissue, confirming its role in stabilizing the epithelial lining of the optic cavity and supporting an epidermal evolutionary origin.

**Supplementary Video 2. Immunolocalization of visual r-opsin (He-R-Opsin-V) in the eye of *H. exemplaris*.** High-resolution confocal stack shows He-R-Opsin-V (turquoise) mainly localized to the rhabdomeric microvilli within the pigment cup. Anterior: up. The pigment cell is visualized via HRP immunolabeling (magenta), confirming its role in optical shielding. No signal is detected in the ciliary photoreceptor or surrounding tissues, confirming He-R-Opsin-V as the sole visual opsin and the eye’s primary photopigment.

**Supplementary Video 3. Triple HCR-FISH of *He-r-opsin-v*, *He-c-opsin-1*, and *He-c-opsin-2* in *H. exemplaris*.** Simultaneous detection reveals *He-r-opsin-v* (magenta) restricted to the rhabdomeric cell of the eye. *He-c-opsin-1* (yellow) is expressed in a dorsomedian cluster outside the brain and co-expressed with *He-c-opsin-2* in anterior and outer lobe brain clusters, as well as in the ciliary photoreceptor cell of the eye. Anterior: up. The expression pattern of *He-c-opsin-1* and *He-c-opsin-2* overlaps in the brain and eye, suggesting shared roles in non-visual light sensing. No signal is detected for *He-r-opsin-v* in the brain or extracerebral regions, confirming its exclusive ocular function.

**Supplementary Video 4. Double HCR-FISH of *He-r-opsin-v* and *He-c-opsin-1* in *H. exemplaris*.** Simultaneous detection reveals *He-r-opsin-v* (magenta) in the rhabdomeric cell and *He-c-opsin-1* (yellow) in the ciliary photoreceptor cell of the eye. Anterior: up. In the brain, *He-c-opsin-1* is expressed in anterior, inner lobe, and outer lobe clusters, with additional labeling in two pairs of ventrolateral clusters outside the brain. *He-r-opsin-v* is absent from the brain, confirming its exclusive ocular function and highlighting functional compartmentalization between visual and non-visual photoreceptive pathways.

**Supplementary Video 5. HCR-FISH of *He****-**c-opsin-2*** **in *H. exemplaris*.** High-resolution confocal stack shows *He-c-opsin-2* (yellow) in the ciliary photoreceptor cell of the eye. Anterior: up. It is additionally expressed in anterior, inner lobe, and outer lobe brain clusters, and in two pairs of ventrolateral clusters outside the brain. The expression pattern mirrors that of *He-c-opsin-1*, suggesting shared roles in non-visual light sensing and neural integration.

**Supplementary Video 6. Double HCR-FISH of *He-r-opsin-v* and *He-c-opsin-3* in *H. exemplaris*.** Simultaneous detection reveals *He-r-opsin-v* (magenta) restricted to the rhabdomeric cell of the eye, with no signal in the brain. Anterior: left. In contrast, *He-c-opsin-3* (yellow) is expressed in a single outer brain lobe neuron and shows no extracerebral expression in the ventrolateral head region—unlike *He-c-opsin-1* and *He-c-opsin-2*. This distinct pattern suggests a specialized, non-visual role in central brain processing.

**Supplementary Video 7. HCR-FISH of *He-r-opsin-nv* in *H. exemplaris*.** High-resolution confocal stack reveals non-visual *He**-**r-opsin-nv* (yellow) expression in inner lobe, posterior cluster within each outer brain lobe and a medial cluster adjacent to each eye. Anterior: up. This spatial separation confirms the non-visual function of *He**-**r-opsin-nv* in central sensory integration.

**Supplementary Video 8. HCR-FISH of *He*-*neuropsin* in *H. exemplaris*.** High-resolution confocal stack reveals *He*-*neuropsin* expression in medial brain regions adjacent to the eye, inner lobes, and posterior outer brain lobes. Anterior: up. No extracerebral signal is detected in the head. The pattern suggests roles in non-visual light sensing, neural integration, or circadian regulation, consistent with its proximity to PDF-positive neurons.

