## Supplementary material for "Extensive opsin gene expansion and non-cerebral origin of the minimalist eye in a model tardigrade": Dutta_et_al_Supplementary_Materials: suppl_fig_2.pdf

K296 NPxxY Tripeptide

NM_001014890.2 *Bos taurus* rhodopsin.............................FGPIFMTIPAFFAKTSAVYNPVIYIMMNKQFR
PX981454 *Hypsibius exemplaris He-r-opsin-v* clone_LH5.........LSPILSQLPALCAKTAAVYNPLVYAISHPKYR
PX981455 *Hypsibius exemplaris He-r-opsin-nv* clone_1SD3......LTPLVAQLPAIFAKTAACYNPVVYVLAHQRYR
PX981456 *Hypsibius exemplaris He-r-opsin-nv* clone_1SD6......LTPLVAQLPAIFAKTAACYNPIVYVFAHQRYR
PX981457 *Hypsibius exemplaris He-r-opsin-nv* clone_1SD8......LTPLVAQLPAIFGKTAACYNPVVYVLAHQRYR
PX981458 *Hypsibius exemplaris He-r-opsin-nv* clone_1SD9......LTPLVAQLPAIFAKTAACYNPVVYVLAHQRYR
PX981459 *Hypsibius exemplaris He-r-opsin-nv* clone_1SD10.....LTPLVAQLPAIFAKTAACYNPVVYVLAHQRYR
PX981460 *Hypsibius exemplaris He-r-opsin-nv* clone_1LH9......LTPLVAQLPAIFAKTAACYNPVVYVLAHQRYR
PX981461 *Hypsibius exemplaris He-r-opsin-nv* clone_1LH10.....LTPLVAQLPAIFGKTAACYNPVVYVLAHQRYR
PX981462 *Hypsibius exemplaris He-r-opsin-nv* clone_1LH12.....LTPLVAQLPAIFGKTAACYNPVVYVLAHQRYR
PX981463 *Hypsibius exemplaris He-r-opsin-nv* clone_1LH13.....LTPLVAQLPAIFAKTAACYNPVVYVLAHQRYR
PX981464 *Hypsibius exemplaris He-r-opsin-nv* clone_1LH14.....LTPLVAQLPAIFAKTAACYNPIVYVFAHQRYR
PX981465 *Hypsibius exemplaris He-r-opsin-nv* clone_1LH17.....LTPLVAQLPAIFAKTAACYNPVVYVLAHQRYR
PX981466 *Hypsibius exemplaris He-r-opsin-nv* clone_1LH18.....LTPLVAQLPAIFGKTAACYNPVVYVLAHQRYR
PX981467 *Hypsibius exemplaris He-r-opsin-nv* clone_1LH19.....LTPLVAQLPAIFAKTAACYNPVVYVLAHQRYR
PX981468 *Hypsibius exemplaris He-r-opsin-nv* clone_1LH21.....LTPLVAQLPAIFAKTAACYNPVVYVLAHQRYR
PX981469 *Hypsibius exemplaris He-r-opsin-nv* clone_1LH22.....LTPLVAQLPAIFGKTAACYNPVVYVLAHQRYR
PX981470 *Hypsibius exemplaris He-r-opsin-nv* clone_1LH23.....LTPLVAQLPAIFAKTAACYNPVVYVLAHQRYR
PX981471 *Hypsibius exemplaris He-r-opsin-nv* clone_1LH24.....LTPLVAQLPAIFAKTAACYNPVVYVLAHQRYR
PX981472 *Hypsibius exemplaris He-r-opsin-nv* clone_2SD10.....LTPLVAQLPAIFAKTAACYNPVVYVLAHQRYR
PX981473 *Hypsibius exemplaris He-r-opsin-nv* clone_2LH3......LTPLVAQLPAIFAKTAACYNPVVYVLAHQRYR
PX981474 *Hypsibius exemplaris He-r-opsin-nv* clone_2LH9......LTPLVAQLPAIFAKTAACYNPVVYVLAHQRYR
PX981475 *Hypsibius exemplaris He-r-opsin-nv* clone_2LH11.....LTPLVAQLPAIFAKTAACYNPVVYVLAHQRYR
*PX981476 Hypsibius exemplaris He-r-opsin-nv clone_2LH12*.....LTPLVAQLPAIFAKTAACYNPVVYVLAHQRYR
*PX981477 Hypsibius exemplaris He-r-opsin-nv clone_2LH13*.....LTPLVAQLPAIFAKTAACYNPVVYVLAHQRYR
*PX981478 Hypsibius exemplaris He-r-opsin-nv clone_2LH15*.....LTPLVAQLPAIFAKTAACYNPVVYVLAHQRYR
PX981479 *Hypsibius exemplaris He-r-opsin-nv* clone_2LH16.....LTPLVAQLPAIFAKTAACYNPVVYVLAHQRYR
PX981480 *Hypsibius exemplaris He-r-opsin-nv* clone_2LH17.....LTPLVAQLPAIFAKTAACYNPVVYVLAHQRYR
PX981481 *Hypsibius exemplaris He-r-opsin-nv* clone_2LH18.....LTPLVAQLPAIFAKTAACYNPVVYVLAHQRYR
PX981482 *Hypsibius exemplaris He-r-opsin-nv* clone_2LH20.....LTPLVAQLPAIFAKTAACYNPVVYVLAHQRYR
PX981483 *Hypsibius exemplaris He-r-opsin-nv* clone_2LH22.....LTPLVAQLPAIFAKTAACYNPVVYVLAHQRYR
PX981484 *Hypsibius exemplaris He-r-opsin-nv* clone_3SD2......LTPLVAQLPAIFGKSAACYNPIVYVFAHQRYR
*PX981485 Hypsibius exemplaris He-r-opsin-nv clone_3SD8*......LTPLVAQLPAIFGKSAACYNPIVYVFAHQRYR
*PX981486 Hypsibius exemplaris He-r-opsin-nv clone_3SD9*......LTPLVAQLPAIFGKSAACYNPIVYVFAHQRYR
*PX981487 Hypsibius exemplaris He-r-opsin-nv clone_3SD10*.....LTPLVAQLPAIFGKSAACYNPIVYVFAHQRYR
*PX981488 Hypsibius exemplaris He-r-opsin-nv clone_3LH11*.....LTPLVAQLPAIFGKSAACYNPIVYVFAHQRYR
*PX981489 Hypsibius exemplaris He-r-opsin-nv clone_3LH12*.....LTPLVAQLPAIFGKSAACYNPIVYVFAHQRYR
*PX981490 Hypsibius exemplaris He-r-opsin-nv clone_3LH17*.....LTPLVAQLPAIFGKSAACYNPIVYVFAHQRYR
*PX981491 Hypsibius exemplaris He-r-opsin-nv clone_3LH19*.....LTPLVAQLPAIFGKSAACYNPIVYVFAHQRYR
*PX981492 Hypsibius exemplaris He-r-opsin-nv clone_3LH21*.....LTPLVAQLPAIFGKSAACYNPIVYVFAHQRYR
PX981493 *Hypsibius exemplaris* He-c-opsin-1 clone_1LH5.........LTATATAVPAIFAKSSIVYNPIIYAFMNVQLR
PX981494 *Hypsibius exemplaris* He-c-opsin-2 clone_2LH7.........VTPAAATAPAVLAKSSIIYNPIIYVFMNPQIR
PX981495 *Hypsibius exemplaris* He-c-opsin-3 clone_2LH5.........ITAPVATLPAIFAKSSIIYNPIIYAIMNPQFR

Suppl_fig_2
