## Supplementary material for "Extensive opsin gene expansion and non-cerebral origin of the minimalist eye in a model tardigrade": Dutta_et_al_Supplementary_Materials: suppl_fig_3.pdf

```
                                                                                                     86 90 91
NM_001014890.2 Bos taurus rhodopsin............................ INFLTLYVTVQHKKLRTPLNYILLNLAVADLFMVFGGFTTTLYTSLHG
AY377391.1 Homo sapiens OPN5.................................... GNGYVLYMSSRRKKKLRPAEIMTINLAVCDLGISVVGKPFTIISCFCH
PX981496 Hypsibius exemplaris He-neuropsin clone_cN1........ ANGVILVGLGRQRRRLTALDILMGNLAVTNVMLALVTYPMTVVSLAFS
PX981497 Hypsibius exemplaris He-neuropsin clone_cN3........ ANGVILVGLSRQRRRLIALDILMGNLAVTNTMLALVTYPMTVASLAFS
PX981498 Hypsibius exemplaris He-neuropsin clone_cN6........ ANGVILVGLGRQRRRLTALDILMGNLAVTNVMLALVTYPMTVVSLAFS
PX981499 Hypsibius exemplaris He-neuropsin clone_cN7........ ANGVILVGLGRQRRRLTALDILMGNLAVTNVMLALVTYPMTVVSLAFS
PX981500 Hypsibius exemplaris He-neuropsin clone_cN9........ ANGVILVGLSRQRRRLIALDILMGNLAVTNTMLALVTYPMTVASLAFS
PX981501 Hypsibius exemplaris He-neuropsin clone_cN10....... ANGVILVGLSRQRRRLIALDILMGNLAVTNTMLALVTYPMTVASLAFS
PX981502 Hypsibius exemplaris He-neuropsin clone_cN13....... ANGVILVGLSRQRRRLIALDILMGNLAVTNTMLALVTYPMTVASLAFS
PX981503 Hypsibius exemplaris He-neuropsin clone_cN14....... ANGVILVGLSRQRRRLIALDILMGNLAVTNTMLALVTYPMTVASLAFS
PX981504 Hypsibius exemplaris He-neuropsin clone_cN15....... ANGVILVGLSRQRRRLIALDILMGNLAVTNTMLALVTYPMTVASLAFS
PX981505 Hypsibius exemplaris He-neuropsin clone_cN16....... ANGVILVGLSRQRRRLIALDILMGNLAVTNTMLALVTYPMTVASLAFS
PX981506 Hypsibius exemplaris He-neuropsin clone_cN17....... ANGVILVGLGRQRRRLTALDILMGNLAVTNVMLALVTYPMTVVSLAFS
PX981507 Hypsibius exemplaris He-neuropsin clone_cN18....... ANGVILVGLSRQRRRLIALDILMGNLAVTNTMLALVTYPMTVASLAFS
PX981508 Hypsibius exemplaris He-neuropsin clone_cN19....... ANGVILVGLSRQRRRLIALDILMGNLAVTNTMLALVTYPMTVASLAFS
PX981510 Hypsibius exemplaris He-neuropsin clone_cN22....... ANGVILVGLSRQRRRLIALDILMGNLAVTNTMLALVTYPMTVASLAFS
PX981511 Hypsibius exemplaris He-neuropsin clone_cN23....... ANGVILVGLGRQRRRLTALDILMGNLAVTNVMLALVTYPMTVVSLAFS
PX981512 Hypsibius exemplaris He-neuropsin clone_cN25....... ANGVILVGLSRQRRRLIALDILMGNLAVTNTMLALVTYPMTVASLAFS
PX981513 Hypsibius exemplaris He-neuropsin clone_cN26....... ANGVILVGLSRQRRRLIALDILMGNLAVTNTMLALVTYPMTVASLAFS
PX981514 Hypsibius exemplaris He-neuropsin clone_cN27....... ANGVILVGLSRQRRRLIALDILMGNLAVTNTMLALVTYPMTVASLAFS
PX981515 Hypsibius exemplaris He-neuropsin clone_cN28....... ANGVILVGLSRQRRRLIALDILMGNLAVTNTMLALVTYPMTVASLAFS
PX981516 Hypsibius exemplaris He-neuropsin clone_cN29....... ANGVILVGLSRQRRRLIALDILMGNLAVTNVMLALVTYPMTVVSLAFS
PX981517 Hypsibius exemplaris He-neuropsin clone_cN30....... ANGVILVGLSRQRRRLIALDILMGNLAVTNTMLALVTYPMTVASLAFS
PX981518 Hypsibius exemplaris He-neuropsin clone_cN34....... ANGVILVGLGRQRRRLTALDILMGNLAVTNVMLALVTYPMTVVSLAFS
PX981519 Hypsibius exemplaris He-neuropsin clone_cN38....... ANGVILVGLSRQRRRLIALDILMGNLAVTNTMLALVTYPMTVASLAFS
PX981520 Hypsibius exemplaris He-neuropsin clone_cN45....... ANGVILVGLSRQRRRLIALDILMGNLAVTNTMLALVTYPMTVASLAFS
PX981521 Hypsibius exemplaris He-neuropsin clone_cN48....... ANGVILVGLSRQRRRLIALDILMGNLAVTNTMLALVTYPMTVASLAFS
PX981522 Hypsibius exemplaris He-neuropsin clone_cN54....... ANGVILVGLSRQRRRLIALDILMGNLAVTNTMLALVTYPMTVASLAFS
PX981523 Hypsibius exemplaris He-neuropsin clone_cN55....... ANGVILVGLSRQRRRLIALDILMGNLAVTNTMLALVTYPMTVASLAFS
PX981525 Hypsibius exemplaris He-neuropsin clone_cN61....... ANGVILVGLSRQRRRLIALDILMGNLAVTNTMLALVTYPMTVASLAFS
PX981526 Hypsibius exemplaris He-neuropsin clone_cN62....... ANGVILVGLGRQRRRLTALDILMGNLAVTNVMLALVTYPMTVVSLAFS
PX981527 Hypsibius exemplaris He-neuropsin clone_cN64....... ANGVILVGLSRQRRRLIALDILMGNLAVTNTMLALVTYPMTVASLAFS
PX981528 Hypsibius exemplaris He-neuropsin clone_cN65....... ANGVILVGLGRQRRRLTALDILMGNLAVTNVMLALVTYPMTVVSLAFS
PX981529 Hypsibius exemplaris He-neuropsin clone_cN66....... ANGVILVGLSRQRRRLIALDILMGNLAVTNTMLALVTYPMTVASLAFS
PX981530 Hypsibius exemplaris He-neuropsin clone_cN67....... ANGVILVGLSRQRRRLIALDILMGNLAVTNTMLALVTYPMTVASLAFS
PX981531 Hypsibius exemplaris He-neuropsin clone_cN68....... ANGVILVGLSRQRRRLIALDILMGNLAVTNTMLALVTYPMTVASLAFS
PX981532 Hypsibius exemplaris He-neuropsin clone_cN70....... ANGVILVGLGRQRRRLTALDILMGNLAVTNTMLALVTYPMTVASLAFS
PX981533 Hypsibius exemplaris He-neuropsin clone_cN71....... ANGVILVGLSRQRRRLIALDILMGNLAVTNTMLALVTYPMTVASLAFS
PX981534 Hypsibius exemplaris He-neuropsin clone_cN72....... ANGVILVGLSRQRRRLIALDILMGNLAVTNTMLALVTYPMTVASLAFS
PX981535 Hypsibius exemplaris He-neuropsin clone_cN73....... ANGVILVGLSRQRRRLIALDILMGNLAVTNTMLALVTYPMTVASLAFS
PX981536 Hypsibius exemplaris He-neuropsin clone_cN74....... ANGVILVGLSRQRRRLIALDILMGNLAVTNTMLALVTYPMTVASLAFS
PX981537 Hypsibius exemplaris He-neuropsin clone_cN75....... ANGVILVGLSRQRRRLIALDILMGNLAVTNTMLALVTYPMTVASLAFS
PX981538 Hypsibius exemplaris He-neuropsin clone_cN78....... ANGVILVGLGRRRRRLTALDILMGNLAVTNVMLALVTYPMTVVSLAFS
PX981539 Hypsibius exemplaris He-neuropsin clone_cN79....... ANGVILVGLSRQRRRLIALDILMGNLAVTNTMLALVTYPMTVASLAFS
PX981540 Hypsibius exemplaris He-neuropsin clone_cN82....... ANGVILVGLGRQRRRLTALDILMGNLAVTNVMLALVTYPMTVVSLAFS
PX981541 Hypsibius exemplaris He-neuropsin clone_cN83....... ANGVILVGLSRQRRRLIALDILMGNLAVTNTMLALVTYPMTVASLAFS
PX981542 Hypsibius exemplaris He-neuropsin clone_cN96....... ANGVILVGLGRQRRRLTALDILMGNLAVTNVMLALVTYPMTVVSLAFS
PX981543 Hypsibius exemplaris He-neuropsin clone_cN99....... ANGVILVGLSRQRRRLIALDILMGNLAVTNTMLALVTYPMTVASLAFS
PX981544 Hypsibius exemplaris He-neuropsin clone_cN115...... ANGVILVGLGRQRRRLTALDILMGNLAVTNVMLALVTYPMTVVSLAFS
PX981545 Hypsibius exemplaris He-neuropsin clone_cN123...... ANGVILVGLGRQRRRLTALDILMGNLAVTNVMLALVTYPMTVVSLAFS
PX981546 Hypsibius exemplaris He-neuropsin clone_cN127...... ANGVILVGLSRQRRRLIALDILMGNLAVTNTMLALVTYPMTVASLAFS
PX981547 Hypsibius exemplaris He-neuropsin clone_cN143...... ANGVILVGLSRQRRRLIALDILMGNLAVTNTMLALVTYPMTVASLAFS
PX981548 Hypsibius exemplaris He-neuropsin clone_cN154...... ANGVILVGLGRQRRRLTALDILMGNLAVTNTMLALVTYPMTVASLAFS
PX981549 Hypsibius exemplaris He-neuropsin clone_cN160...... ANGVILVGLSRQRRRLIALDILMGNLAVTNTMLALVTYPMTVASLAFS
PX981550 Hypsibius exemplaris He-neuropsin clone_cN164...... ANGVILVGLSRQRRRLIALDILMGNLAVTNTMLALVTYPMTVASLAFS
PX981551 Hypsibius exemplaris He-neuropsin clone_cN170...... ANGVILVGLGRQRRRLTALDILMGNLAVTNVMLALVTYPMTVVSLAFS
PX981552 Hypsibius exemplaris He-neuropsin clone_cN173...... ANGVILVGLSRQRRRLIALDILMGNLAVTNTMLALVTYPMTVASLAFS
PX981553 Hypsibius exemplaris He-neuropsin clone_cN176...... ANGVILVGLGRQRRRLTALDILMGNLAVTNVMLALVTYPMTVVSLAFS
PX981554 Hypsibius exemplaris He-neuropsin clone_cN178...... ANGVILVGLSRQRRRLIALDILMGNLAVTNTMLALVTYPMTVASLAFS
PX981555 Hypsibius exemplaris He-neuropsin clone_cN180...... ANGVILVGLSRQRRRLIALDILMGNLAVTNTMLALVTYPMTVASLAFS
MTYJ01000071_He-Neuropsin_CDS.............................. ANGVILVGLSRQRRRLIALDILMGNLAVTNTMLALVTYPMTVASLAFS
MTYJ01000117_He-Neuropsin_CDS.............................. ANGVILVGLGRQRRRLTALDILMGNLAVTNVMLALVTYPMTVVSLAFS
PZ005554 Hypsibius exemplaris He-neuropsin-a1 clone_gN4..... ANGVILVGLGRQRRRLTALDILMGNLAVTNVMLALVTYPMTVVSLAFS
PZ005555 Hypsibius exemplaris He-neuropsin-a2 clone_gN5..... ANGVILVGLSRQRRRLIALDILMGNLAVTNTMLALVTYPMTVASLAFS
PZ005556 Hypsibius exemplaris He-neuropsin-a3 clone_gN7..... ANGVILVGLSRQRRRLIALDILMGNLAVTNTMLALVTYPMTVASLAFS
PZ005557 Hypsibius exemplaris He-neuropsin-a4 clone_gN9..... ANGVILVGLSRQRRRLIALDILMGNLAVTNTMLALVTYPMTVASLAFS
PZ005558 Hypsibius exemplaris He-neuropsin-a5 clone_gN13.... ANGVILVGLSRQRRRLIALDILMGNLAVTNTMLALVTYPMTVASLAFS
PZ005559 Hypsibius exemplaris He-neuropsin-a6 clone_gN15.... ANGVILVGLSRQRRRLIALDILMGNLAVTNTMLALVTYPMTVASLAFS
PZ005560 Hypsibius exemplaris He-neuropsin-a7 clone_gN17.... ANGVILVGLSRQRRRLIALDILMGNLAVTNTMLALVTYPMTVASLAFS
PZ005561 Hypsibius exemplaris He-neuropsin-a8 clone_gN24.... ANGVILVGLSRQRRRLIALDILMGNLAVTNTMLALVTYPMTVASLAFS
PZ005562 Hypsibius exemplaris He-neuropsin-a9 clone_gN30.... ANGVILVGLSRQRRRLIALDILMGNLAVTNTMLALVTYPMTVASLAFS
PZ005563 Hypsibius exemplaris He-neuropsin-b1 clone_gN2..... ANGVILVGLGRQRRRLTALDILMGNLAVTNVMLALVTYPMTVVSLAFS
PZ005564 Hypsibius exemplaris He-neuropsin-b2 clone_gN14.... ANGVILVGLGRQRRRLTALDILMGNLAVTNVMLALVTYPMTVVSLAFS
PZ005565 Hypsibius exemplaris He-neuropsin-b3 clone_gN18.... ANGVILVGLGRQRRRLTALDILMGNLAVTNVMLALVTYPMTVVSLAFS
PZ005566 Hypsibius exemplaris He-neuropsin-b4 clone_gN20.... ANGVILVGLGRQRRRLTALDILMGNLAVTNVMLALVTYPMTVVSLAFS
PZ005567 Hypsibius exemplaris He-neuropsin-b5 clone_gN21.... ANGVILVGLGRQRRRLTALDILMGNLAVTNVMLALVTYPMTVVSLAFS
PZ005568 Hypsibius exemplaris He-neuropsin-b6 clone_gN25.... ANGVILVGLGRQRRRLTALDILMGNLAVTNVMLALVTYPMTVVSLAFS
PZ005569 Hypsibius exemplaris He-neuropsin-b7 clone_gN27.... ANGVILVGLGRQRRRLTALDILMGNLAVTNVMLALVTYPMTVVSLAFS
PZ005570 Hypsibius exemplaris He-neuropsin-b8 clone_gN29.... ANGVILVGLGRQRRRLTALDILMGNLAVTNVMLALVTYPMTVVSLAFS
PZ005571 Hypsibius exemplaris He-neuropsin-b9 clone_gN31.... ANGVILVGLGRQRRRLTALDILMGNLAVTNVMLALVTYPMTVVSLAFS
```

Suppl_fig_3
