## Supplementary material for "Extensive opsin gene expansion and non-cerebral origin of the minimalist eye in a model tardigrade": Dutta_et_al_Supplementary_Materials: suppl_table_1.docx

**Supplementary Table 1. Revised nomenclature for opsins in *H. exemplaris*.**

| **Revised nomenclature** | **Hering & Mayer 2014** | **Fleming et al. 2021** | **Gene identifier*** | **Scaffold*** | **Accession*** |
| --- | --- | --- | --- | --- | --- |
| *He-r-opsin-v* | *Hd-r-opsin* | *R-opsin II1* | BV898_10375 | scaffold0088 | MTYJ01000088.1 |
| *He-r-opsin-nv-a1* | *-* | *R-opsin III1* | BV898_16528 | scaffold0250 | MTYJ01000250.1 |
| *He-r-opsin-nv-a2* | *-* | *R-opsin III2* | BV898_16527 | scaffold0250 | MTYJ01000250.1 |
| *He-r-opsin-nv-b1* | *-* | *R-opsin III3* | BV898_16526 | scaffold0250 | MTYJ01000250.1 |
| *He-c-opsin-1* | *Hd-c-opsin1* | *C-opsin 1* | BV898_18200 | scaffold0345 | MTYJ01000345.1 |
| *He-c-opsin-2* | *Hd-c-opsin2* | *C-opsin 2* | BV898_11633 | scaffold0109 | MTYJ01000109.1 |
| *He-c-opsin-3* | *Hd-c-opsin3* | *C-opsin 3* | BV898_11632 | scaffold0109 | MTYJ01000109.1 |
| *He-neuropsin-a* | *Hd-neuropsin* | *Neuropsin 1* | BV898_09176 | scaffold0071 | MTYJ01000071.1 |
| *He-neuropsin-b* | *-* | *Neuropsin 2* | BV898_12046 | scaffold0117 | MTYJ01000117.1 |

* Genome assembly from Yoshida et al. (2017)

Yoshida, Y., Koutsovoulos, G., Laetsch, D.R., et al. (2017). Comparative genomics of the tardigrades *Hypsibius dujardini* and *Ramazzottius varieornatus*. *PLOS Biology* 15:e2002266.
