## Supplementary material for "Extensive opsin gene expansion and non-cerebral origin of the minimalist eye in a model tardigrade": Dutta_et_al_Supplementary_Materials: suppl_table_2.docx

**Supplementary Table 2. Gene-specific primers for amplification of *H. exemplaris* opsin transcripts.**

| **Gene** | **Fragment length** | **Direction** | **Primer sequence (5’🡪3’)** |
| --- | --- | --- | --- |
| *He-r-opsin-v* | 1,533 bp | Forward | ATGTCTGGCGAATTGTACGAAGA |
|  |  | Reverse | TTACACGGTGGTAACTTTTTCCTG |
| *He-r-opsin-nv-a1* | 1,161 bp | Forward | ATGGCCGTCAACACCACC |
|  |  | Reverse | TCAACCGTTCCGTATCTTGACT |
| *He-r-opsin-nv-a2* | 1,323 bp | Forward | ATGGCGCTCAACACCACC |
|  |  | Reverse | TTACGCACAGTAGATGAGGAGCA |
| *He-r-opsin-nv-b1* | 1,089 bp | Forward | ATGACCTTCAACGCCACTATCACC |
|  |  | Reverse | TCATTGACGTTGATGTGGACGYTT |
| *He-c-opsin-1* | 1,152 bp | Forward | ATGACTCTGGAGCAGCATCTCAA |
|  |  | Reverse | TCAAACAGCACTGCTGGGGA |
| *He-c-opsin-2* | 1,275 bp | Forward | ATGACAGACATGGGTAAACTGCACG |
|  |  | Reverse | TTAGGCCGTGAAGGCGTAAGTCC |
| *He-c-opsin-3* | 1,122 bp | Forward | ATGCATGAACACTGGACCGC |
|  |  | Reverse | TTAAAACTGAGACTCAGTAACGCCG |
| *He-neuropsin-a/-b** | 1,221/1,218 bp | Forward | ATGGATCAAAACAGWASCGGGAC |
|  |  | Reverse | TCACGTTGYTTGMTGATTKTTGC |

*Due to sequence similarity degenerated primers were used to amplify *He-neuropsin-a* and *He-neuropsin-b* simultaneously and separated by subsequent cloning.
