## Supplementary material for "Extensive opsin gene expansion and non-cerebral origin of the minimalist eye in a model tardigrade": Dutta_et_al_Supplementary_Materials: suppl_table_3.docx

| **Supplementary Table 3. Polyclonal antibodies against *H. exemplaris* opsins.** | | | | | | | | | | | |
| --- | --- | --- | --- | --- | --- | --- | --- | --- | --- | --- | --- |
| **Marker** | | **Peptide Sequence** | | | **Molecular weight (kDa)** | | | | **Host animal** | **Working concentration (µg/mL)** | |
| He-R-Opsin-V | | | | CIKSPDEEIKGTSSVGKSTRIATGNGSS | | | 2.81 | guinea pig | | | 8 |
| He-C-Opsin-1 | | | | GFSNLQSGPSSAGHKSDSA | | | 1.83 | rabbit | | | 8 |
| He-Neuropsin* | -a/  -b | | CLRYIDQVVNE**S**E**G**PVRSQSG  CLRYIDQVVNE**P**E**R**PVRSQSG | | | 2.34  2.45 | | guinea pig | | | 8–20 |

*Note that due to the similarity of peptide sequences, we cannot exclude the possibility that the single anti-neuropsin antibody used in this study binds both neuropsins.
