## Supplementary material for "Extensive opsin gene expansion and non-cerebral origin of the minimalist eye in a model tardigrade": Dutta_et_al_Supplementary_Materials: suppl_table_4.docx

| **Supplementary Table 4. Secondary antibodies used immunohistochemistry and peptide competition assay.** | | | | |
| --- | --- | --- | --- | --- |
| **Antibody** | **Host animal** | **Fluorophore** | **Concentration** | **Manufacturer** |
| Anti-guinea pig IgG (H+L) | donkey | Alexa Fluor^®^ 488 | 4 µg/mL | Dianova GmbH, Hamburg, Germany |
| Anti-rabbit IgG (H+L) | goat | Alexa Fluor^®^ 488 | 4 µg/mL | Thermo Fisher Scientific Inc., Waltham, MA, USA |
| Anti-rabbit IgG (H+L) | goat | Alexa Fluor^®^ 568 | 4 µg/mL |  |
