## Supplementary material for "Extensive opsin gene expansion and non-cerebral origin of the minimalist eye in a model tardigrade": Dutta_et_al_Supplementary_Materials: Suppl_table_5.docx

| **Target gene** | **Accession No./locus_tag** | **Length**  **(nt)** | **Probe set size** | **Amplifiers** | **Probe working concentration** | **company** |
| --- | --- | --- | --- | --- | --- | --- |
| *He-r-opsin-v* | MTYJ01000088  BV898_10375 | 1,533 | 18 | B1 | 4 µl to 500 µl hybridization buffer | Molecular Instruments, USA |
| *He-r-opsin-nv-a1* | MTYJ01000250  BV898_16528 | 1,161 | 16 | B3 | 10 µl to 500 µl hybridization buffer |  |
| *He-r-opsin-nv-a2* | MTYJ01000250  BV898_16527 | 1,323 | 15 | B1 | 13 µl to 500 µl hybridization buffer |  |
| *He-r-opsin-nv-b1* | MTYJ01000250  BV898_16526 | 1,089 | 15 | B2 | 13 µl to 500 µl hybridization buffer |  |
| *He-c-opsin-1* | MTYJ01000345  BV898_18200 | 1,152 | 8 | B4 | 13 µl to 500 µl hybridization buffer |  |
| *He-c-opsin-2* | MTYJ01000109  BV898_11633 | 1,275 | 16 | B2 | 10 µl to 500 µl hybridization buffer |  |
| *He-c-opsin-3* | MTYJ01000109  BV898_11632 | 1,122 | 15 | B3 | 13 µl to 500 µl hybridization buff |  |
| *He-neuropsin-a* | MTYJ01000071  BV898_09176 | 1,221 | 15 | B4 | 13 µl to 500 µl hybridization buff |  |
| *He-neuropsin-b* | MTYJ01000117  BV898_12046 | 1,218 | 15 | B3 | 13 µl to 500 µl hybridization buff |  |

**Supplementary Table 5. HCR-FISH probe and amplifier details.**
