## Supplementary figures and images for "Extensive opsin gene expansion and non-cerebral origin of the minimalist eye in a model tardigrade"

### suppl_fig_1.pdf

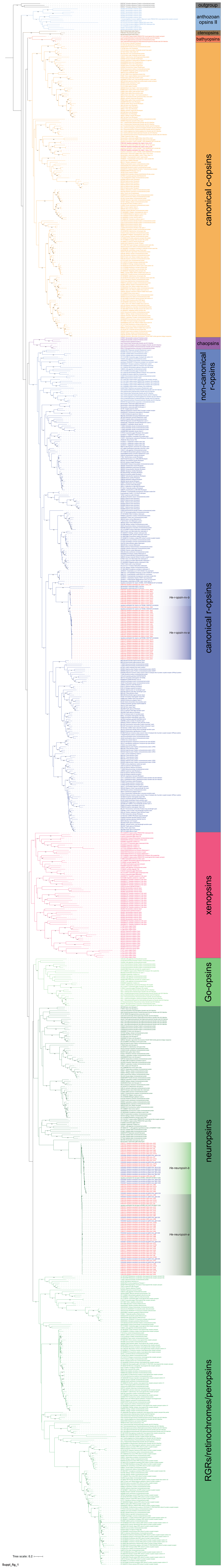

### suppl_fig_4.pdf

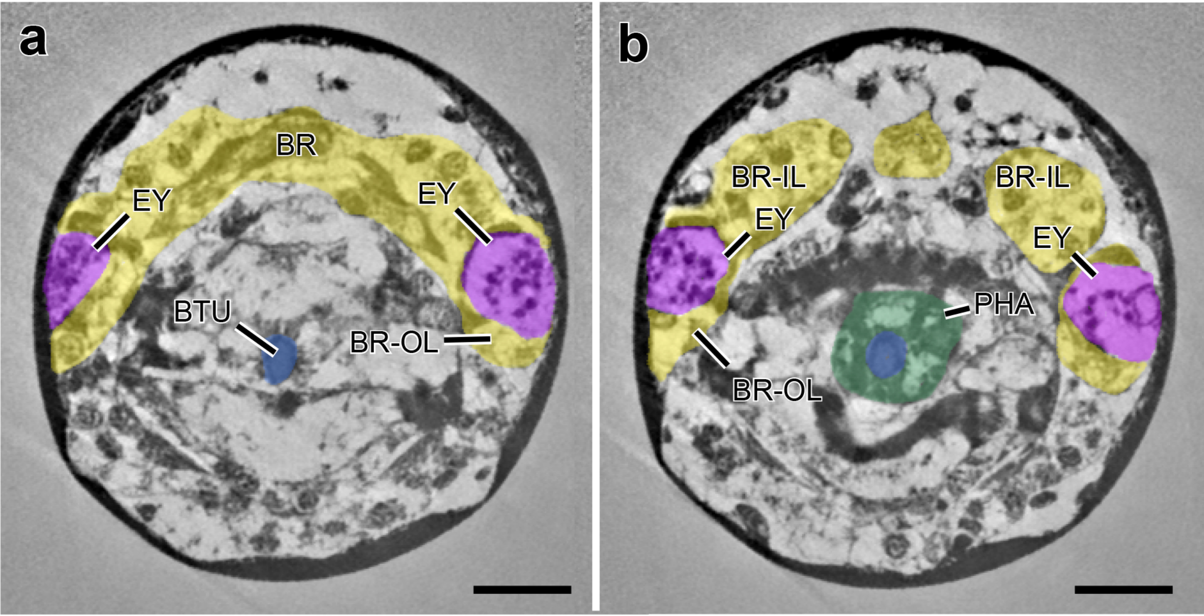


suppl_fig_4

### suppl_fig_5.pdf

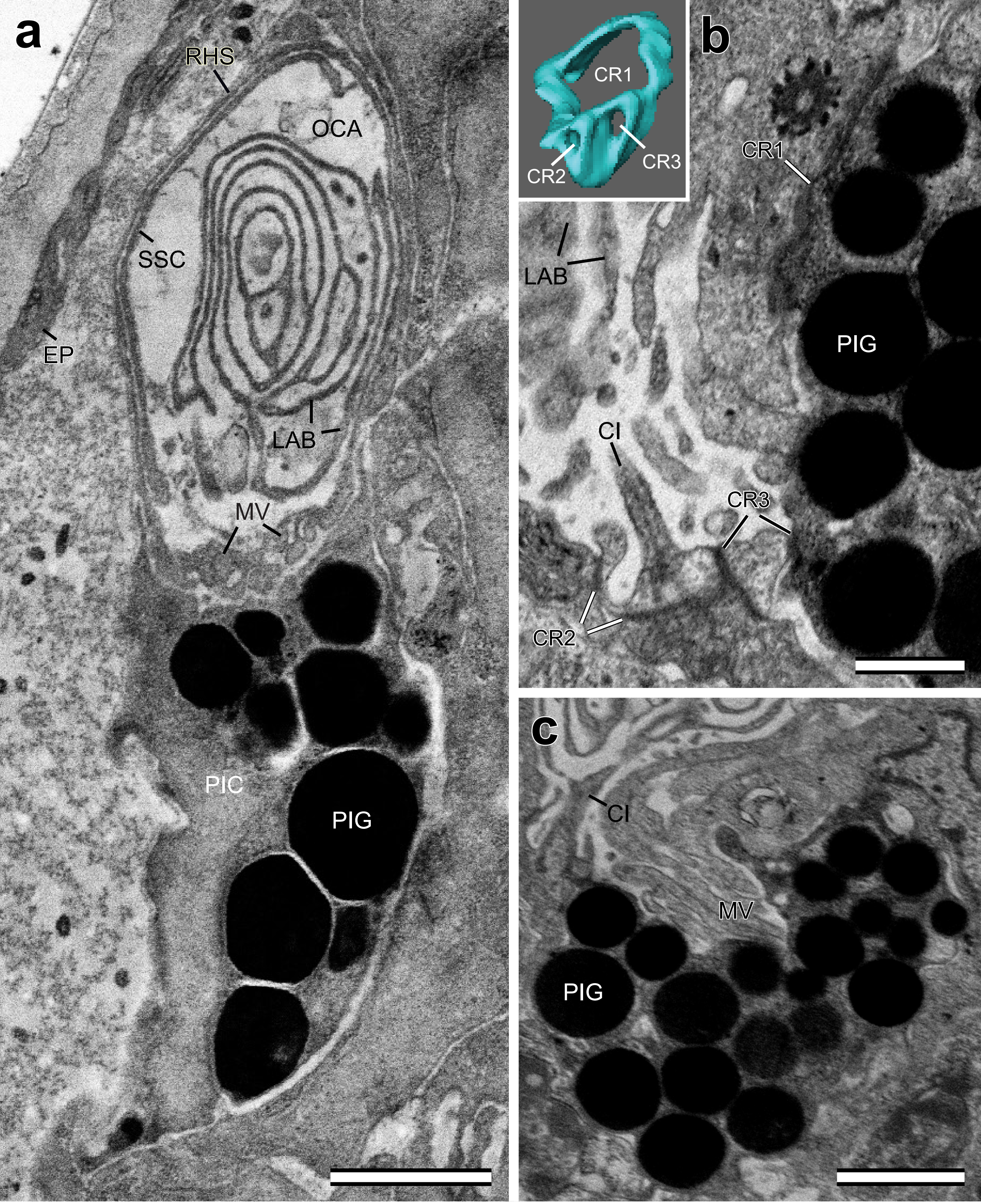


Suppl_fig_5

### suppl_fig_6.pdf

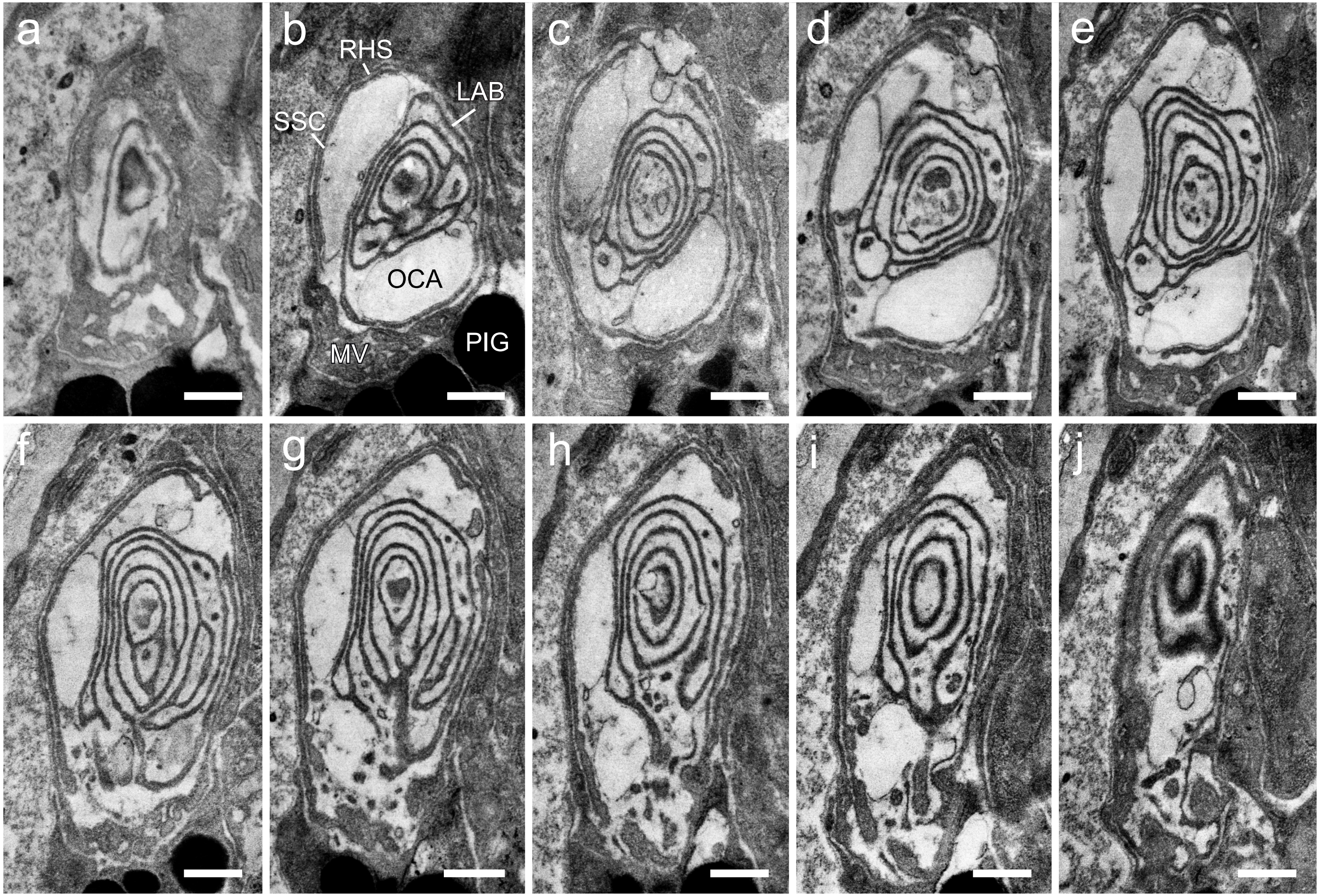


Suppl_fig_6

### suppl_fig_7.pdf

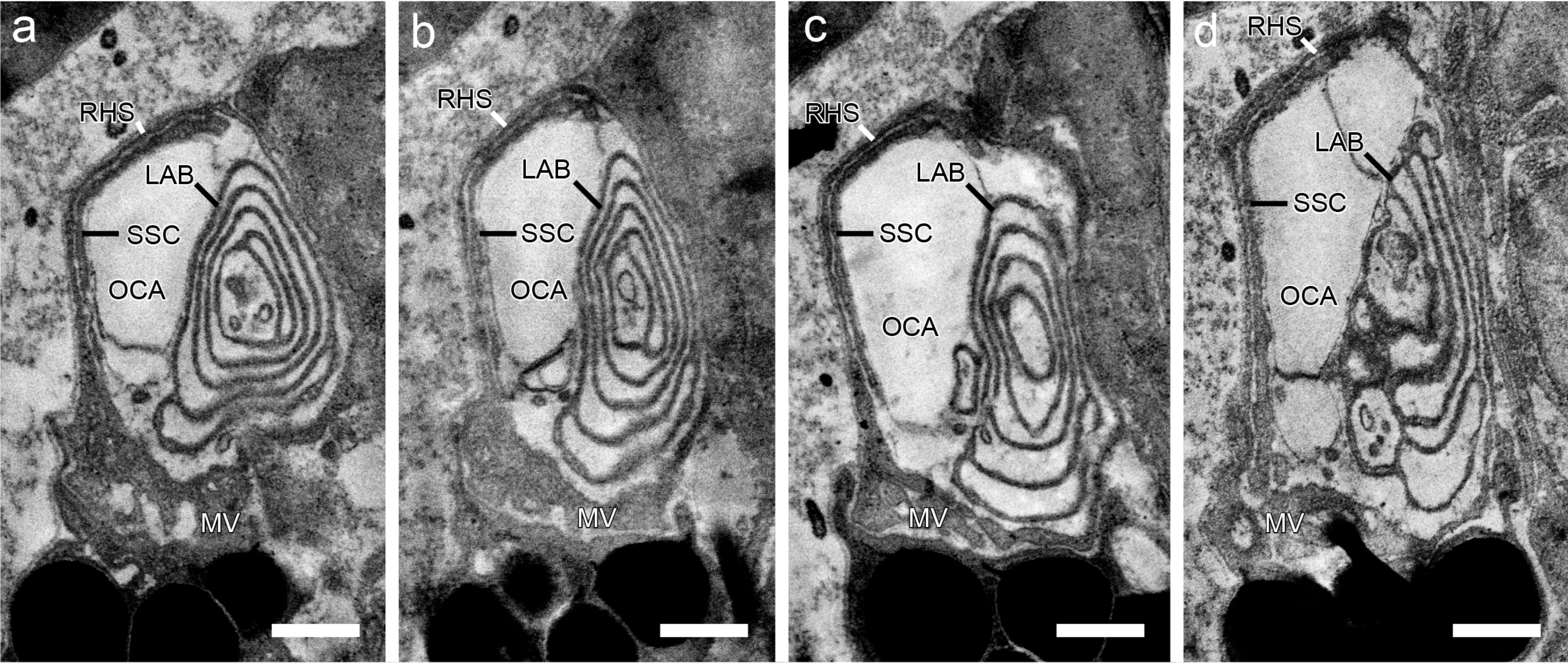

### suppl_fig_8.pdf

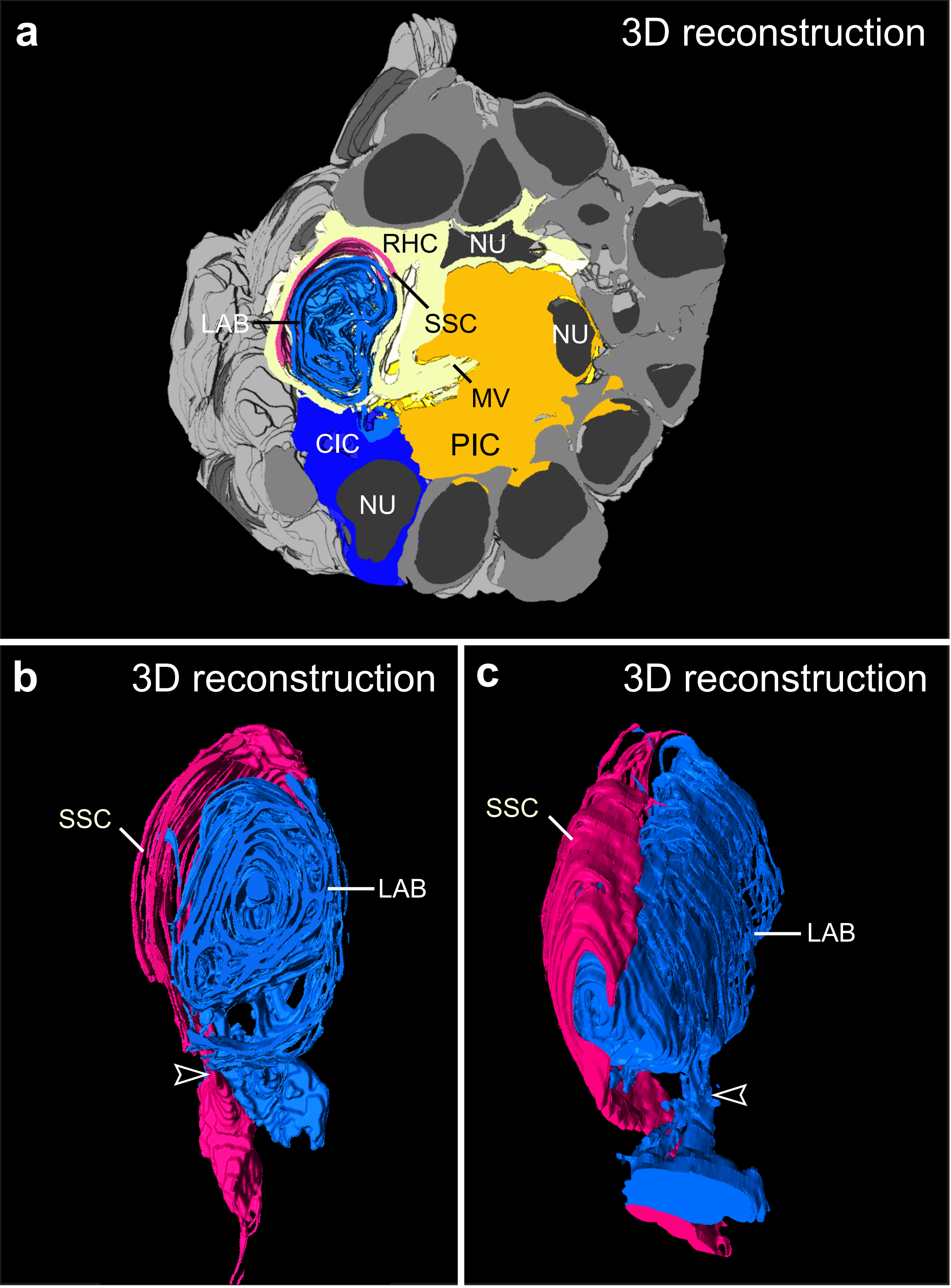

### suppl_fig_9.pdf

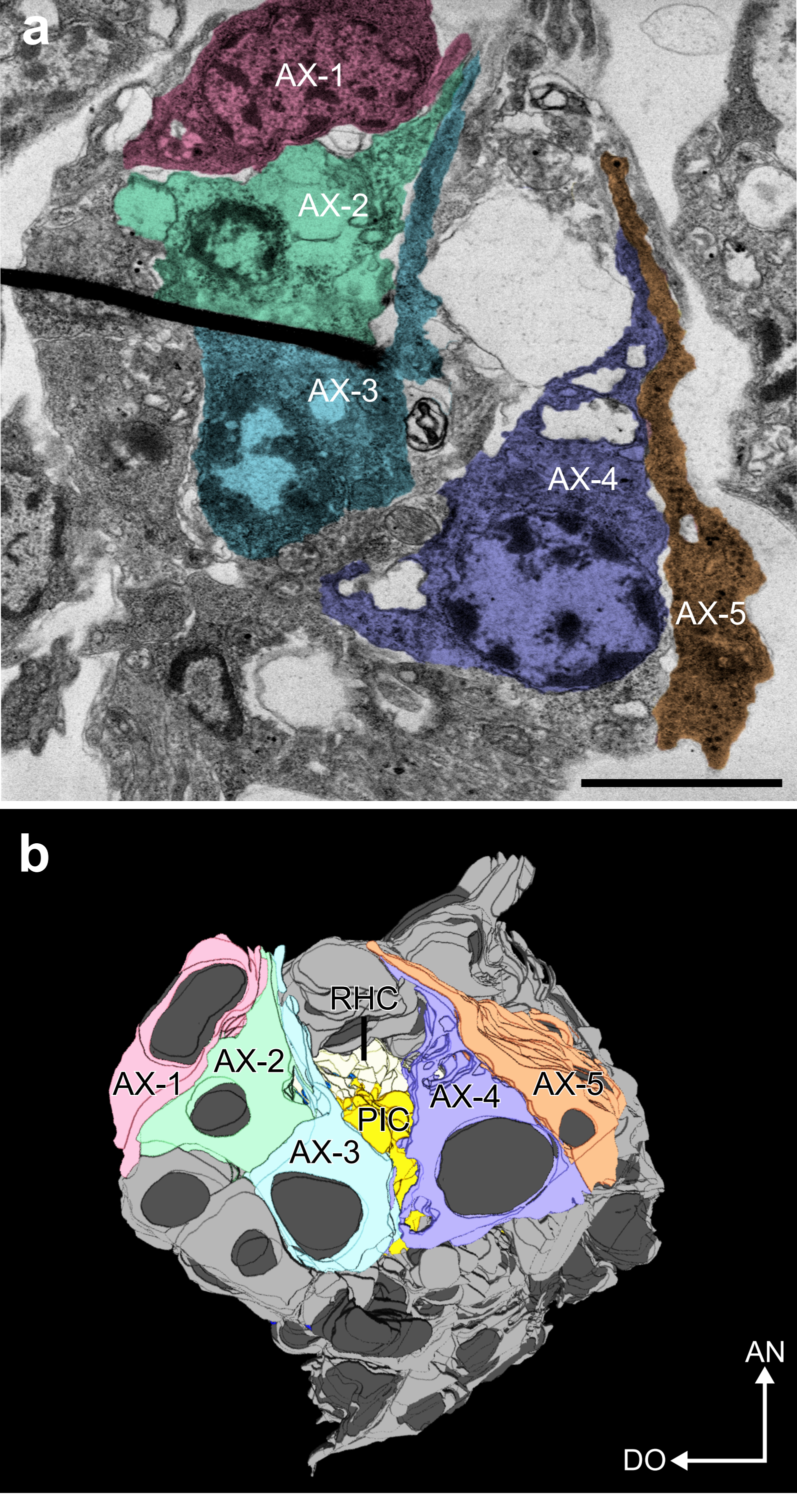


Suppl_fig_9

### suppl_fig_10.pdf

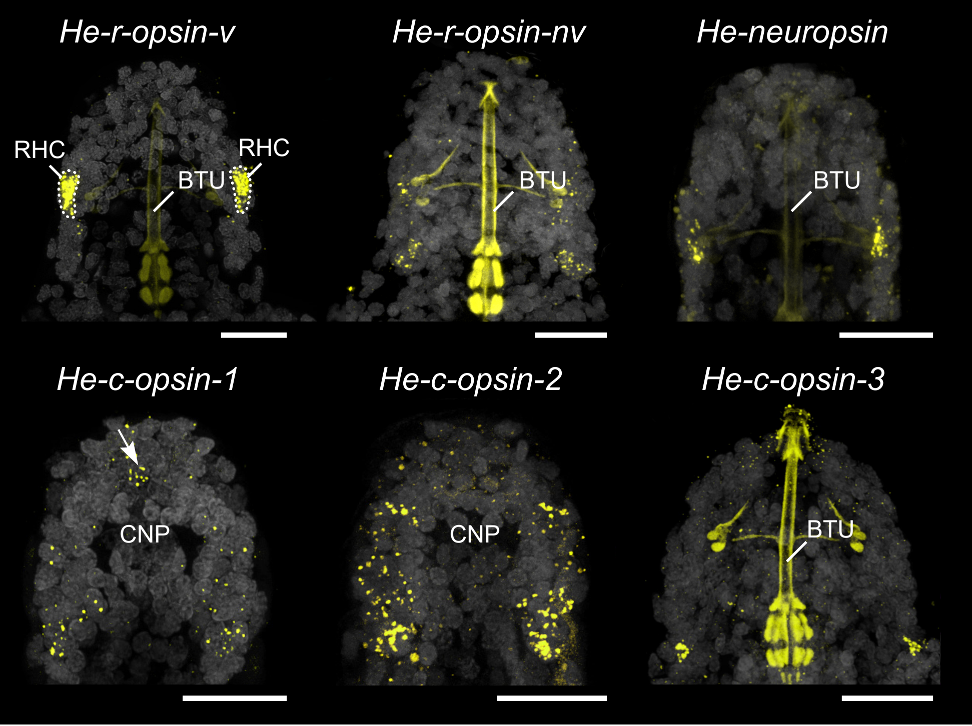

### suppl_fig_11.pdf

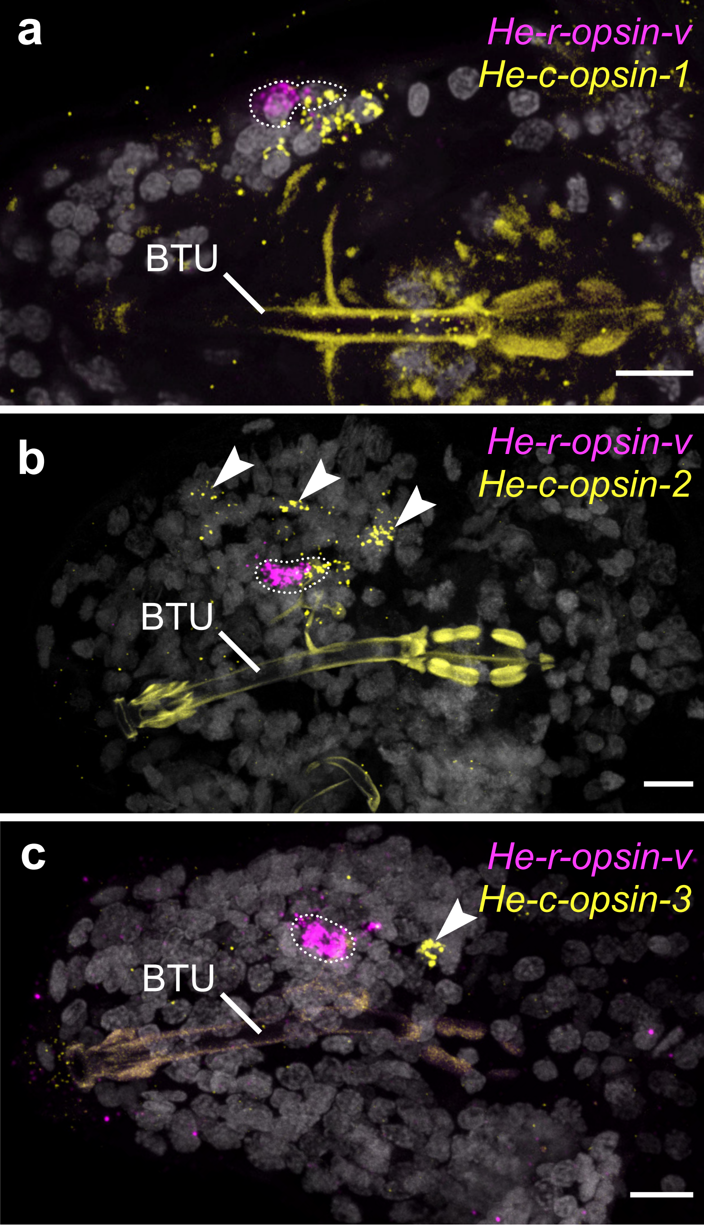

### suppl_fig_12.pdf

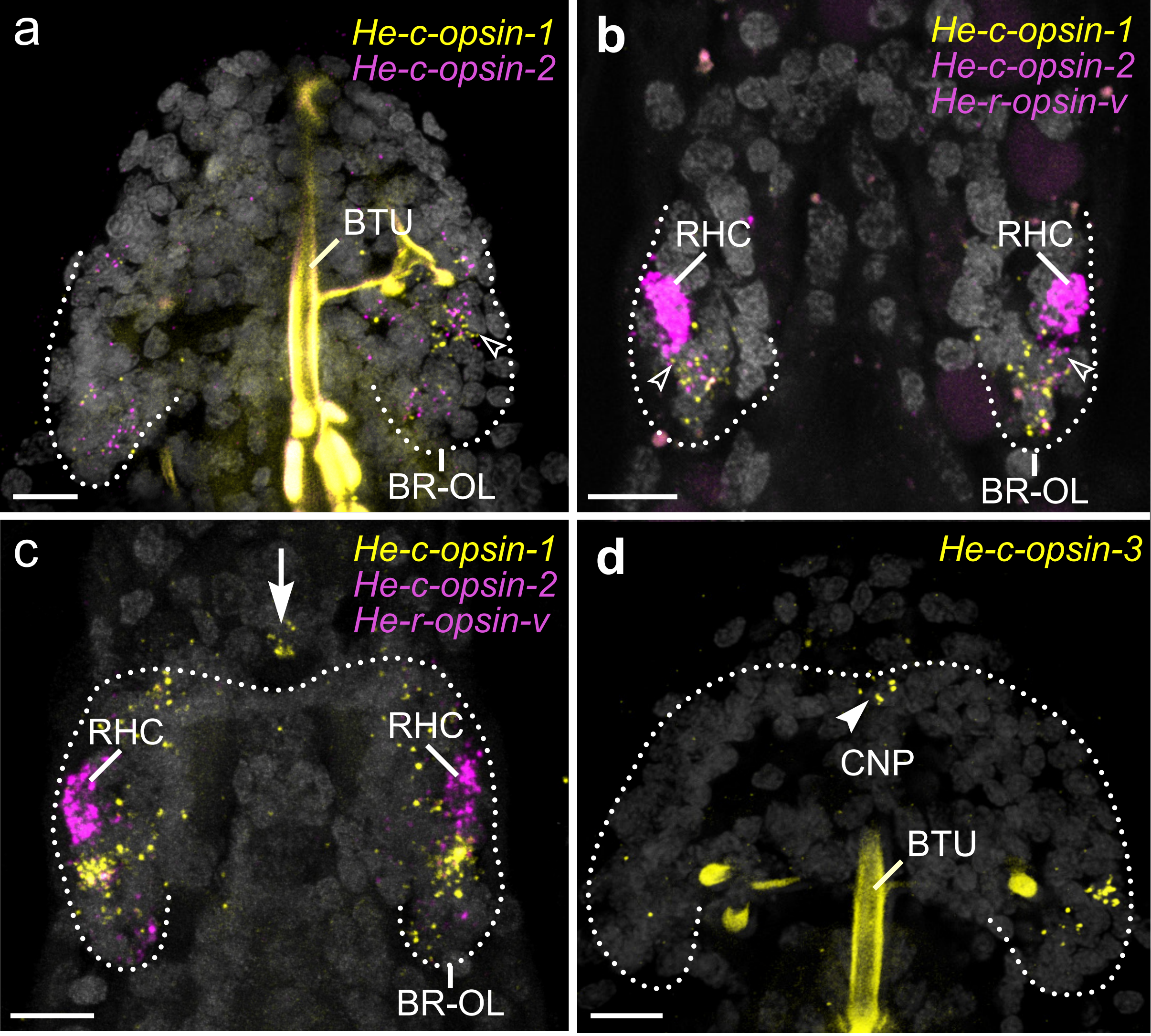

### suppl_fig_13.pdf

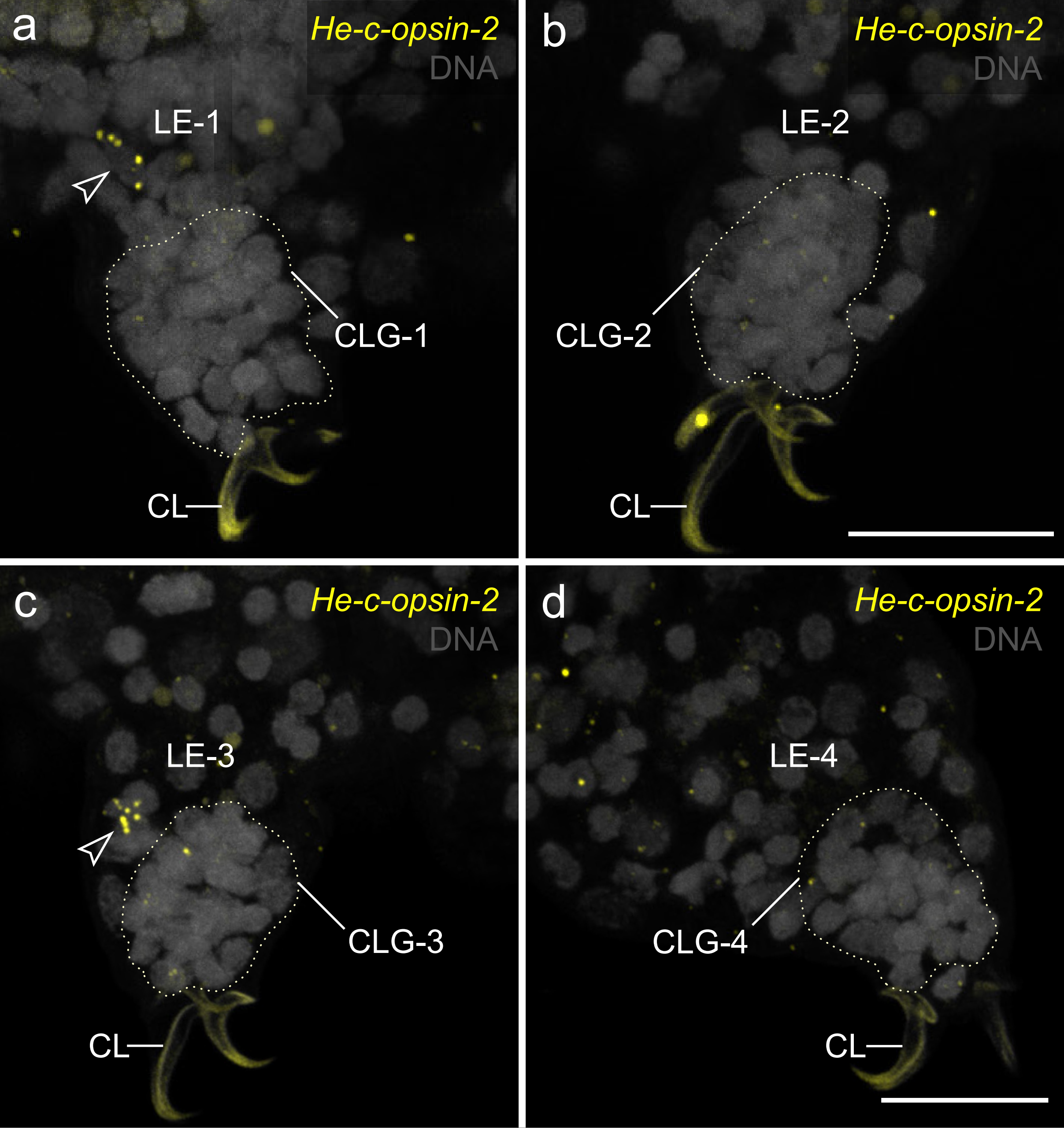

### suppl_fig_14.pdf

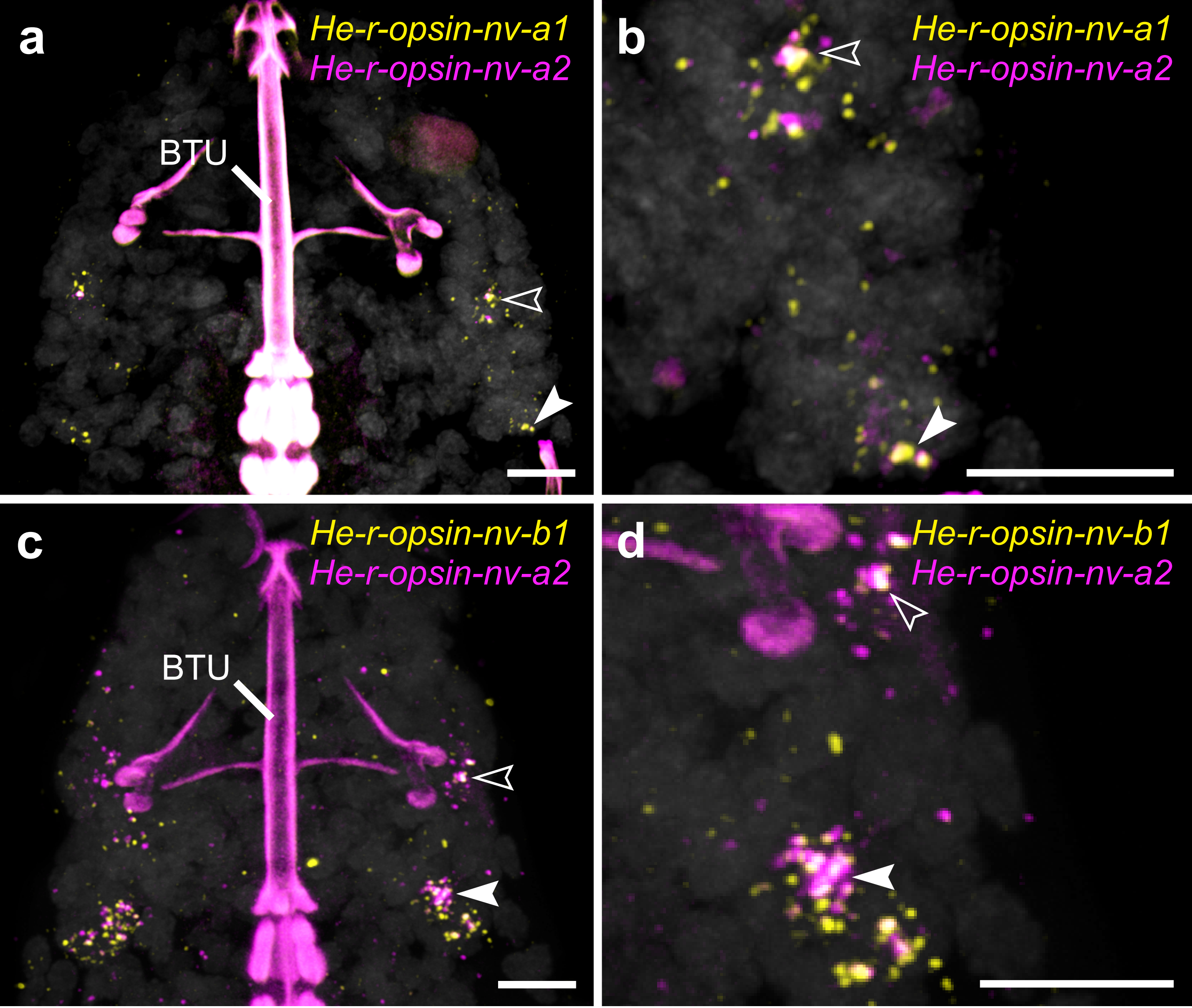

### suppl_fig_15.pdf

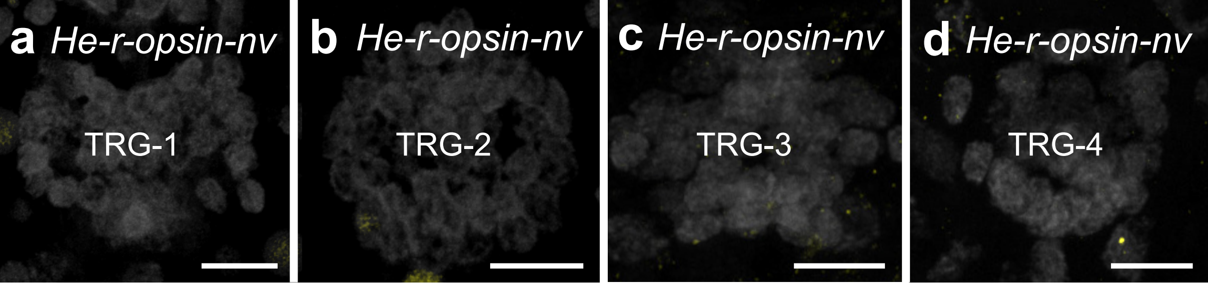

### suppl_fig_16.pdf

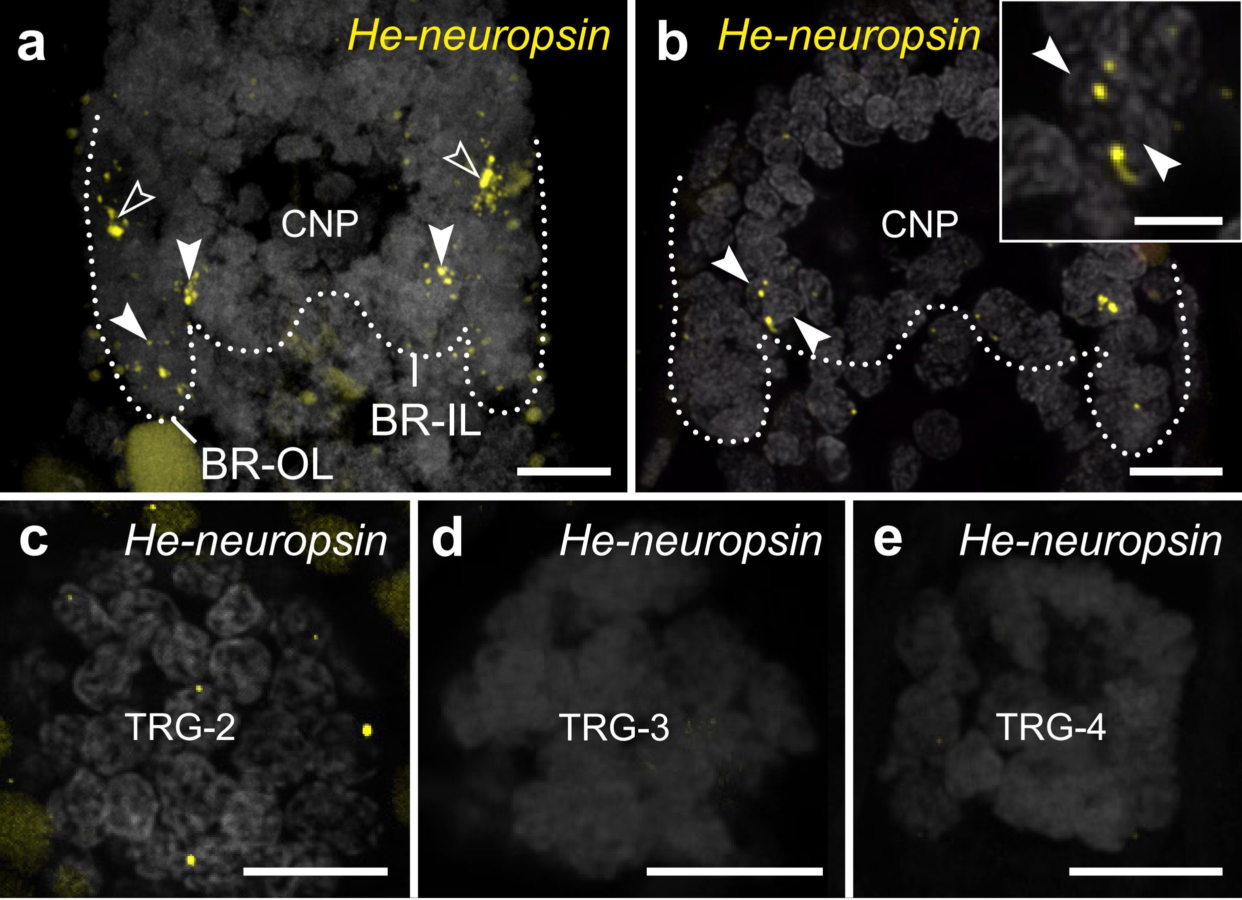

### suppl_fig_17.pdf

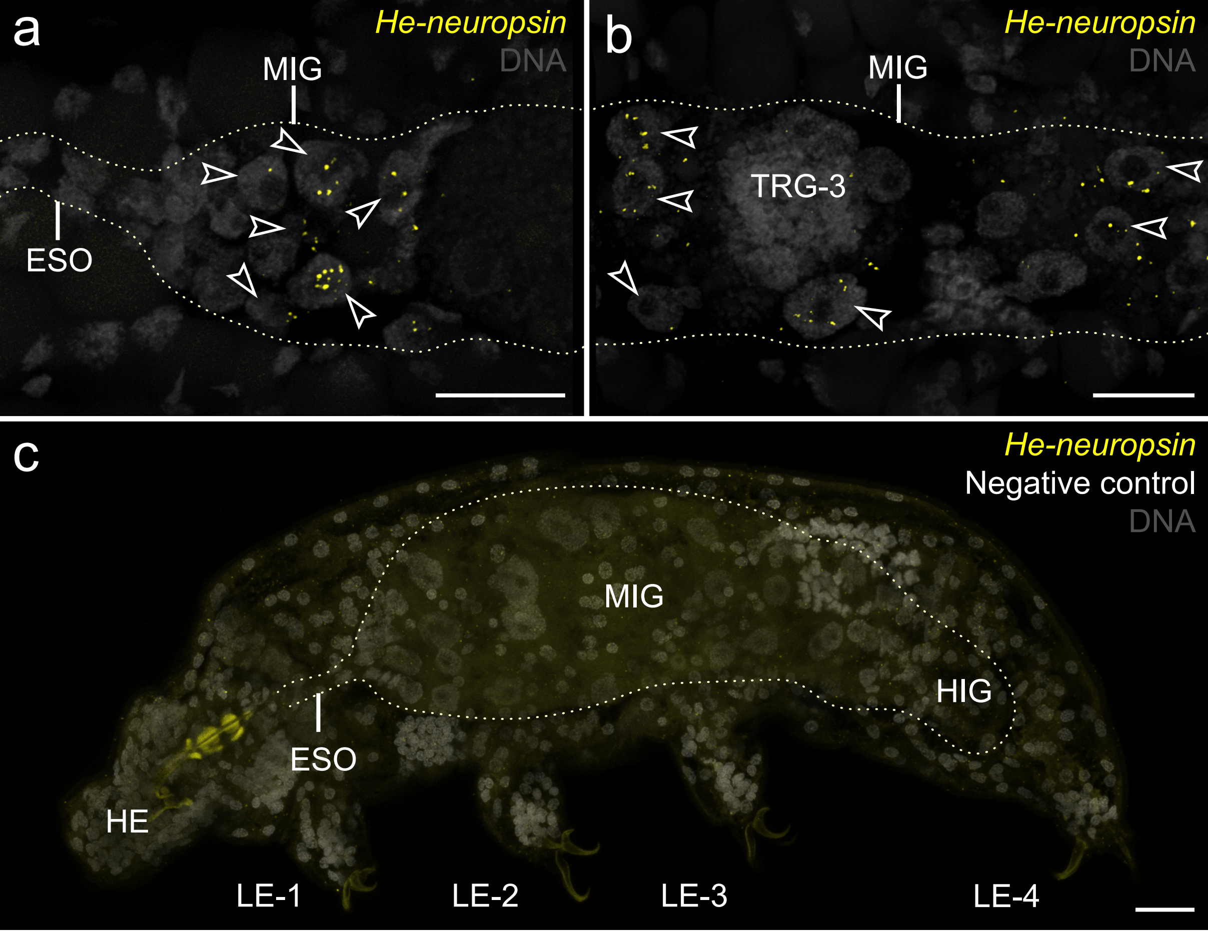

### suppl_fig_18.pdf

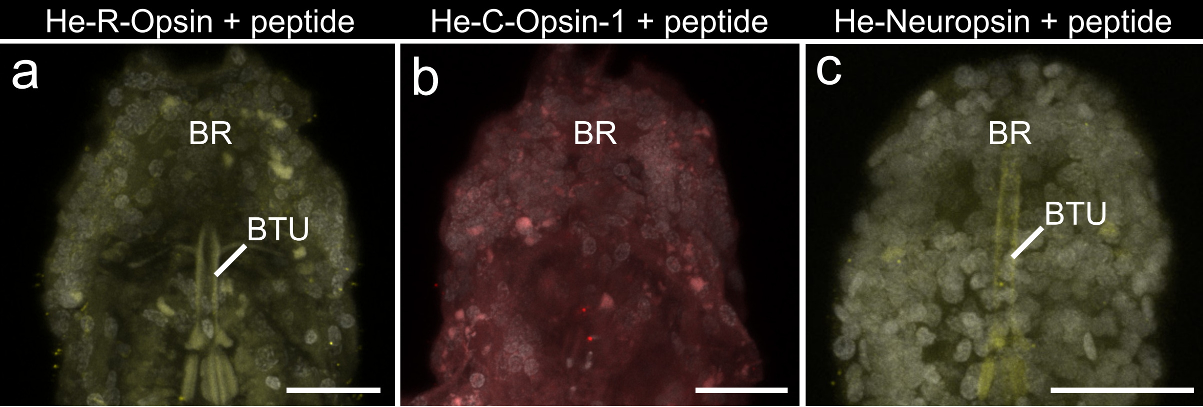
